# Tradeoffs between phage resistance and nitrogen fixation drive the evolution of genes essential for cyanobacterial heterocyst functionality

**DOI:** 10.1101/2023.10.04.560878

**Authors:** Dikla Kolan, Esther Cattan-Tsaushu, Hagay Enav, Zohar Freiman, Nechama Malinsky-Rushansky, Shira Ninio, Sarit Avrani

## Abstract

Harmful blooms caused by diazotrophic (nitrogen-fixing) *cyanobacteria* are becoming increasingly frequent and negatively impact aquatic environments worldwide. Cyanophages (viruses infecting *cyanobacteria*) can potentially regulate cyanobacterial blooms, yet *cyanobacteria* can rapidly acquire mutations that provide protection against phage infection. Here, we provide novel insights into cyanophage:*cyanobacteria* interactions by characterizing the resistance to phages in two species of diazotrophic *cyanobacteria*: *Nostoc* sp. and *Cylindrospermopsis raciborskii*. Our results demonstrate that phage resistance is associated with a fitness tradeoff by which resistant *cyanobacteria* have reduced ability to fix nitrogen and/or to survive nitrogen starvation. Furthermore, we use whole genome sequence analysis of 58 *Nostoc* resistant strains to identify several mutations associated with phage resistance, including in cell surface-related genes, and regulatory genes involved in development and function of heterocysts (cells specialized in nitrogen fixation). Finally, we employ phylogenetic analyses to show that most of these resistance genes are accessory genes whose evolution is impacted by lateral gene transfer events. Together, these results further our understanding of the interplay between diazotrophic *cyanobacteria* and their phages, and suggest that a tradeoff between phage resistance and nitrogen fixation affects the evolution of cell surface-related genes and of genes involved in heterocyst differentiation and nitrogen fixation.

## Introduction

Nitrogen is essential for the vitality of ecosystems. While nitrogen is the most abundant element in the air, atmospheric nitrogen (N_2_) is unavailable for the majority of organisms on earth. By performing nitrogen fixation, diazotrophic microorganisms make it biologically available and thereby represent essential contributors to the global nitrogen budget [1–3]. As nitrogenase, the enzyme that catalyzes nitrogen fixation, is sensitive to oxygen [4], nitrogen fixation predominantly takes place in anaerobic environments. However, diazotrophic *cyanobacteria* represent an exception due to their ability to fix nitrogen under aerobic conditions.

Since oxygen is a byproduct of photosynthesis, *cyanobacteria* have evolved a few strategies to enable nitrogen fixation to occur in parallel with photosynthesis. One example is a spatial separation between the two processes, evolved in some of the filamentous *cyanobacteria* species belonging to the order *Nostocales*. In these species, nitrogen fixation is executed in specialized cells called heterocysts [2] while photosynthesis is performed in vegetative cells. Heterocyst cells enable the microoxic environment needed for the nitrogen fixation reaction by minimizing O_2_ diffusion into the cell via changes to the cell envelope, and can also deplete O_2_ via the use of respiration [5]. The cell envelope of the heterocysts is thicker than that of vegetative cells and consists of two additional external layers: the inner layer (called the heterocyst-specific glycolipid layer, or HGL) reduces the diffusion of gases, including atmospheric O_2_; while the outer layer (called the heterocyst envelope polysaccharide layer, or HEP) provides mechanical support to the HGL [6, 7]. The functional importance of these two layers is highlighted by the observation that mutants with aberrant HEP or HGL are unable to fix N_2_ under aerobic conditions (described as a Fox^-^ phenotype) [8, 9].

The ability to fix nitrogen confers a substantial adaptive advantage to diazotrophic *cyanobacteria* over other phytoplankton species (non-diazotrophic *cyanobacteria* and algae) under nitrogen starvation conditions [10]. This advantage allows diazotrophic strains to form harmful blooms [11, 12], which have become more frequent and harmful in the past decades. While various abiotic factors (such as P-depletion or water mixing) are thought to lead to bloom collapse [13–16], the role of biotic factors in this process remains unknown [14].

Cyanophages (viruses infecting *cyanobacteria*) are one potential biotic factor that regulates cyanobacterial blooms. However, while viruses are known to play an important part in the demise of marine algae blooms [17, 18], the role of cyanophages in cyanobacterial bloom collapse is unclear. Cyanophages are abundant in both marine and terrestrial environments, where their interactions with their hosts can be seen by records of cyanophage sequences in CRISPR arrays in their hosts [19–21] and by prophage remnants found in many cyanobacterial genomes [19, 22, 23]. Moreover, *cyanobacteria* encode various defense mechanisms against phages [24], suggesting a long coevolutionary interplay between these hosts and their phages. *Cyanobacteria* have also been shown to become resistant to the infecting phage by acquiring spontaneous mutations [25–30]. In most cases, these mutations affect the composition of the cell surface of the host, thus resulting in reduced adsorption of phage particles to the host cell surface [9, 25, 31, 32]. Resistance of *cyanobacteria* to phage infection often comes at the cost of a reduction in growth rate [25, 29, 33], enhanced infection by other phages [25, 34], decreased buoyancy [35] or reduction in filament length [35, 36]. Such tradeoffs have a significant role in phage-host dynamics, because they restrict the growth of resistant strains [34]. While a reduction in the growth rate of resistant strains was observed, in most cases the physiological cause of this reduction remains unknown. An exception to that are two observations, in which mutations in the diazotrophic strain *Nostoc* PCC sp. strain 7120 (*Nostoc* 7120) conferring resistance to the cyanophages A-1L and A-4L, abolished the ability to fix N_2_ in aerobic environments [9, 37]. While this tradeoff may have a significant effect on bloom dynamics under nitrogen starvation conditions, the mechanisms by which phage resistance impact nitrogen fixation remain predominantly uncharacterized and the broader relevance of these phenomena beyond *Nostoc* is unclear.

To address these questions, here we report a tradeoff between resistance to phages and nitrogen fixation in two heterocystous *cyanobacteria* species: the model *Nostoc* sp. PCC 7120 and the invasive bloom-forming *Cylindrspermopsis raciborskii*. We isolated 58 resistant substrains of *Nostoc* 7120 resistant to the cyanophages A-4L and/or AN-15 and eight substrains of *C. raciborski* resistant to three CrV-like cyanophages, and show that all 17 *Nostoc* 7120 and eight *C. raciborski* resistant substrains analyzed have reduced ability to fix nitrogen and/or to survive under nitrogen starvation (Table 1). By sequencing the 58 resistant *Nostoc* 7120 substrains, we identify 50 mutations specific to mutant strains. Eleven of the mutant genes were associated with phage resistance, as well as with reduced nitrogen fixation and/or limited growth under nitrogen starvation and were not reported previously as related to nitrogen fixation. Additional phylogenetic analyses of these genes provide insights into their evolutionary history, advancing our understanding of the nitrogen fixation process in heterocystous *cyanobacteria* and of how the evolution of this process is impacted by the tradeoff between phage resistance and heterocyst differentiation.

**Table 1:**
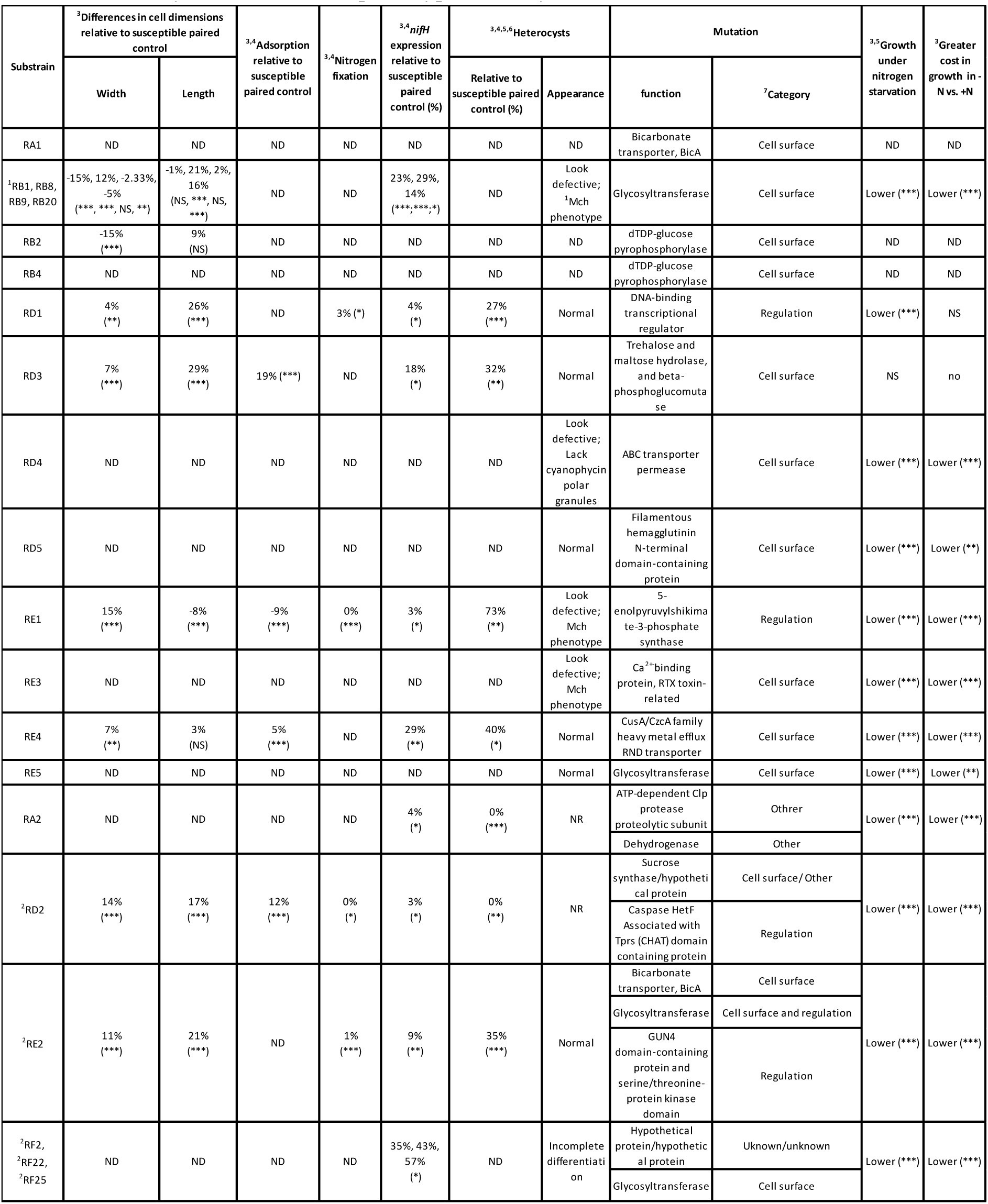
Summary of mutations and phenotypes in analyzed resistant substrains.

## Results and Discussion

### Resistance to phages

To better understand the interactions between diazotrophic *cyanobacteria* and their phages, we isolated 58 substrains of the heterocystous *cyanobacteria Nostoc* PCC 7120 (hereafter *Nostoc* 7120) resistant to the short tail cyanophage Anabaena phage A-4L (hereafter A-4L) and/or the long contractile tail cyanophage AN-15 (hereafter AN-15) (Table S1) [38, 39]. Cell length and width of a subset of these resistant substrains (11 substrains as shown in table 1; see methods for strain selection) was then analyzed and compared to their susceptible paired controls.

The vast majority (10/11) of the resistant substrains analyzed had significant differences in cell width (Fig. 1A,B) and/or length (Fig. 1B, S1) when compared to their susceptible parental strains. Moreover, the cells of one of the resistant substrains (RD2) had an aberrant morphology and were highly granulated (Fig. 1B), which may indicate an accumulation of cell inclusions (such as cyanophycin granules, gas vacuoles, phycobilisomes, carboxysomes, and /or glycogen granules) [40–42]. The observed variation in the phenotypes of the resistant substrains suggests that they carry different mutations.

**Figure 1:**
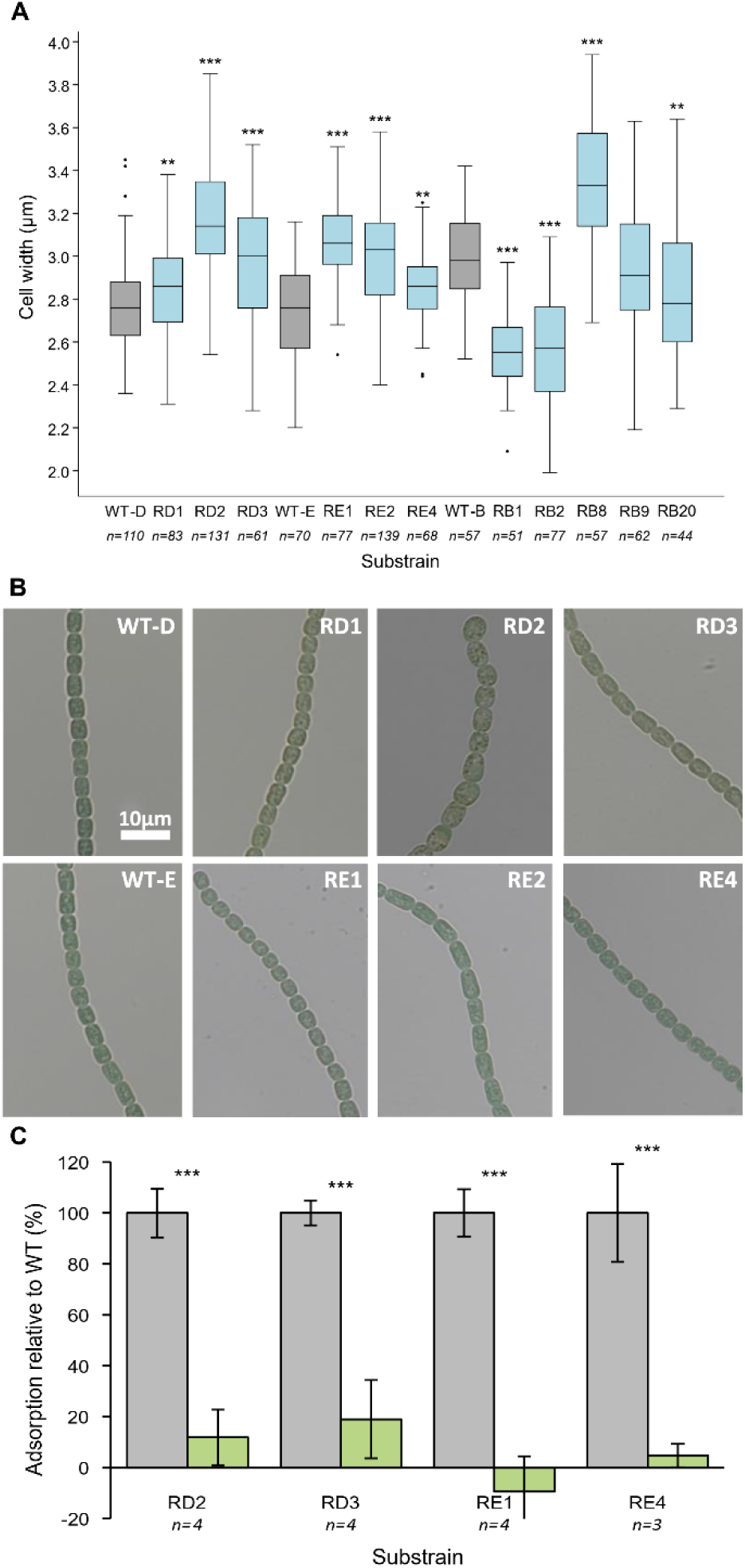
Phenotypes of the resistant substrains of *Nostoc* 7120. Cells width (**A**) of the resistant substrains (light blue) and their susceptible ancestors (WT; grey). Data shown are average +/- standard deviation of 44-139 cells per substrain. **B.** Bright Field images of resistant substrains and their susceptible controls. **C.** Percentage of the adsorbed A-4L phage particles to four resistant substrains of *Nostoc* 7120 (green) relative to the adsorption of the phage to their susceptible paired controls (grey). Data shown are average +/- standard deviation of 3-4 biological replicates. \*\**p*< 0.01 \*\*\**p*< 0.001.

We then examined the mechanisms that enable phage resistance in these strains. The first step in phage infection is the adsorption of the phage to the cell surface of its host [43]. To assess whether resistance was due to impaired attachment, we performed adsorption assays using a lysate of phage A-4L and a subset of 4 of the evolved resistant substrains (see methods for strain selection), as well as their susceptible ancestors. All analyzed resistant substrains displayed a significant reduction in the adsorption of the phage to their cell surface (Fig. 1C, Table 1), suggesting that resistance is due to impaired attachment of the phage to its host, likely as a result of modifications to the bacterial cell surface.

### Cost of resistance

Resistance to phages is often associated with an adaptive cost, manifested many times by a reduction in growth rate [44–47]. To test whether resistance of *Nostoc* 7120 to A-4L and AN-15 affects the growth performance of the host, we compared the growth of a selection of 17 resistant substrains (nine of the 11 substrains analyzed in the previous section as well as eight additional substrains; see table 1) to that of their susceptible ancestor (see methods for strain selection), both under nitrogen replete and nitrogen limitation conditions. Fifteen out of the seventeen examined resistant substrains had a significant reduction in growth compared to controls under nitrogen replete conditions (Fig. 2A, S2, Table 1). Under nitrogen starvation conditions, we observed a significant growth deficiency compared to controls in 16 out of 17 examined substrains (Fig. 2B, S2, Table 1). Moreover, all of these 16 substrains had a significantly greater growth reduction under nitrogen starvation than in nitrogen replete conditions (Fig. 2 and S2, Table 1), with 10 of these being completely unable to grow shortly after nitrogen stepdown (Fig. 2B, S2).

**Figure 2:**
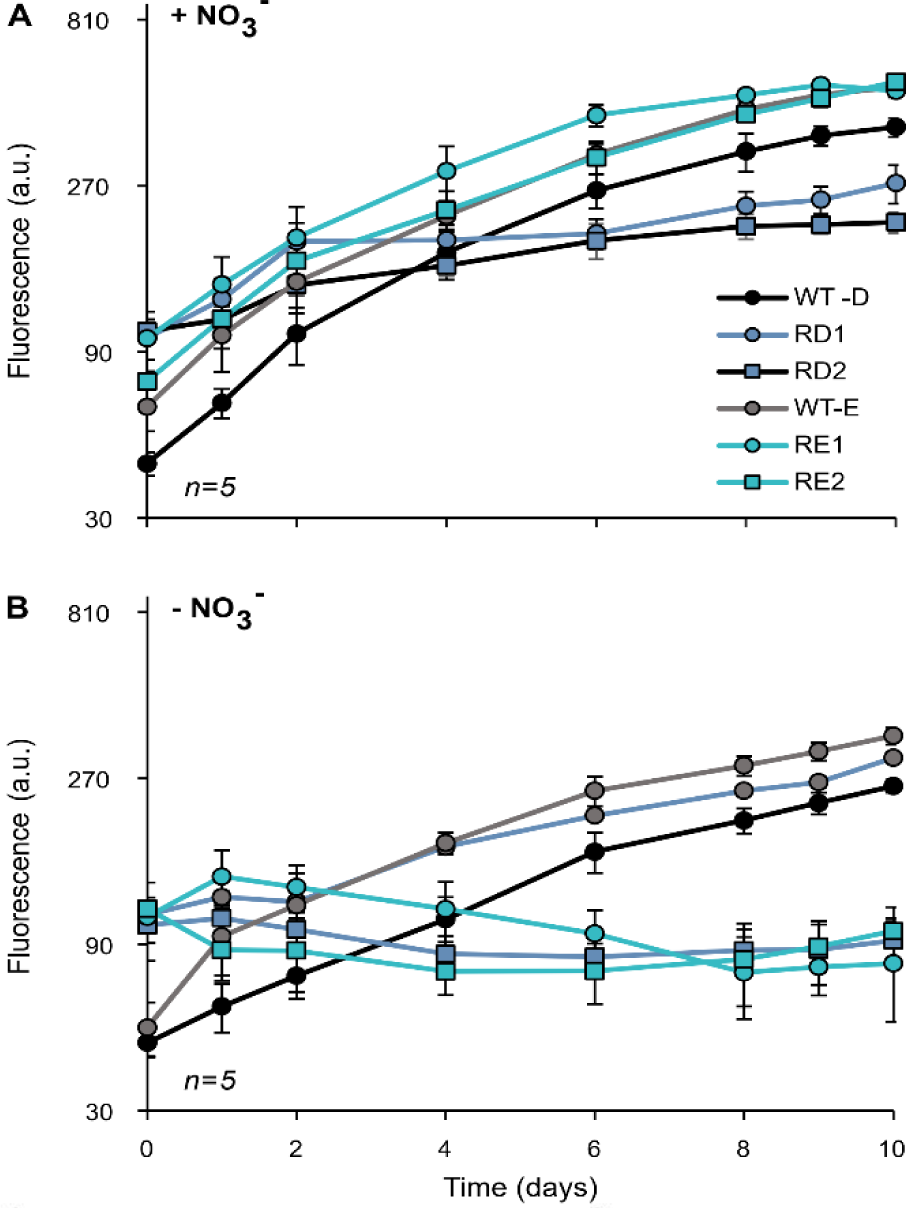
Growth cost in *Nostoc* 7120. Growth dynamics of resistnat substrains of *Nostoc* 7120 and their susceptible controls in nitrogen replete (**A**) and in nitrogen deplete conditions (**B**). The growth of the resistant substrains in nitrogen rich medium was significantly lower than that of the susceptible wild type for RD1, RD2 and RE1 (*p*<0.001). The resistant substrains RD2, RE1, and RE2 had a significantly greater cost under nitrogen starvation than in nitrogen replete medium (*p*<0.001). Data shown are average +/- standard deviation of five biological replicates. Relative cell density is estimated by chlorophyll *a* autofluorescence. a.u: arbitrary units.

### Reduced nitrogen fixation

The reduction in growth of the resistant substrains under nitrogen starvation suggests that phage resistance limits the ability to these *cyanobacteria* to fix nitrogen. To directly test this, we compared the nitrogenase activity of 4 *Nostoc* resistant substrains (see methods for strain selection) to that of the susceptible control strains at 48 hours post nitrogen stepdown. Indeed, all analyzed resistant substrains (4/4) show a complete or nearly complete loss of nitrogenase activity (Fig. 3A, Table 1).

**Figure 3:**
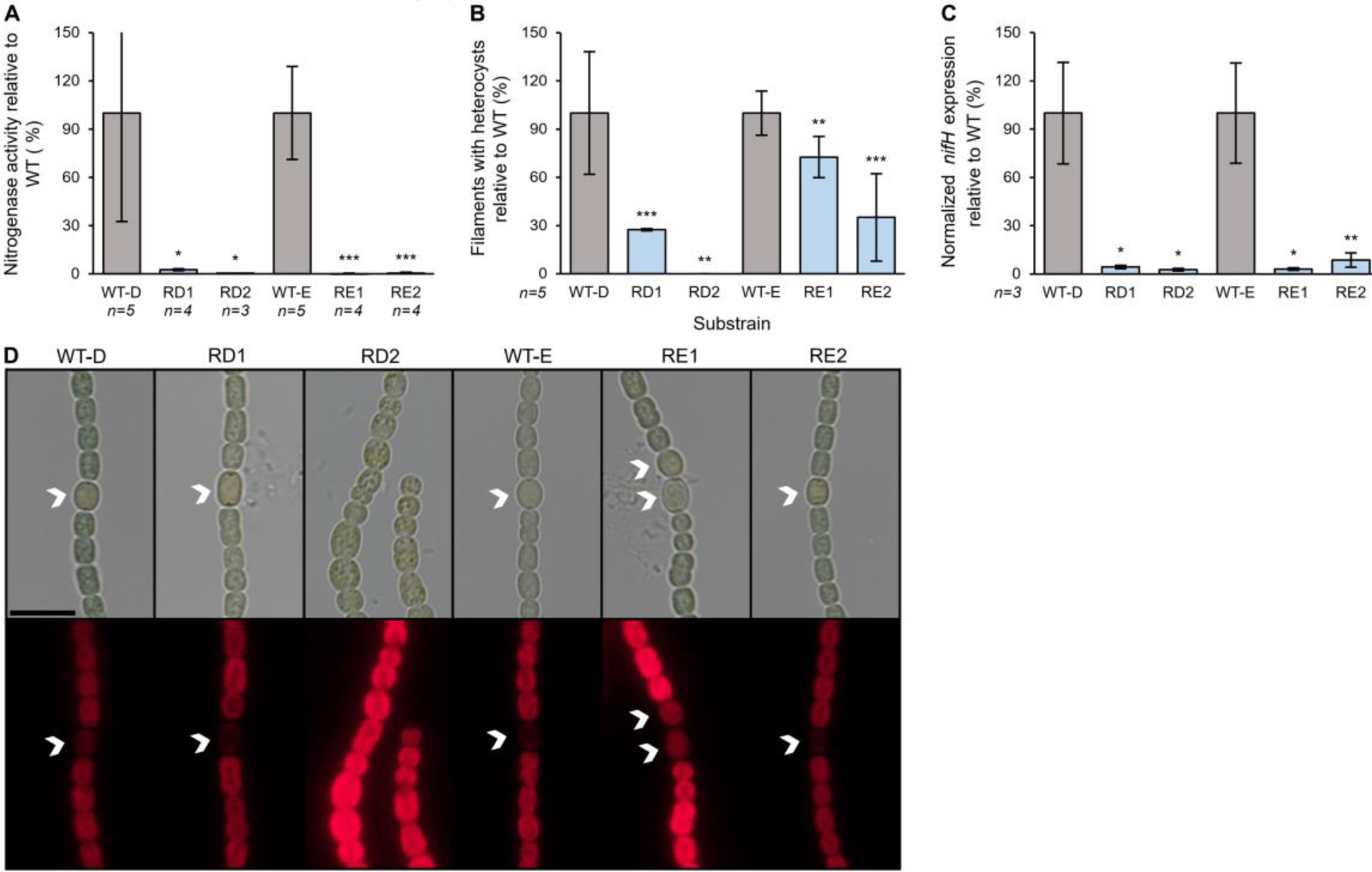
Nitrogen fixation related cost in *Nostoc* 7120. **A.** Nitrogenase activity, relative to the susceptible wild type (grey), of the resistant substrains (light blue) 48 hours after nitrogen stepdown. **B.** Percentage of filaments with heterocysts of the resistant substrains (light blue), relative to the susceptible wild type (grey) 48 hours after nitrogen stepdown. **C.** Expression of *nifH* gene in the resistant substrains (light blue) relative to the susceptible ancestor (grey) 48 hours after nitrogen stepdown. The transcript levels of *nifH* values are normalized to the transcript levels of *rnpB*. Data shown are average +/- standard deviation of 3-5 biological replicates. * *p*< 0.05, ** *p*< 0.01, *** *p*< 0.001. **D.** Bright field images (upper panels) and the corresponding fluorescence images (lower panels; red color represents auto fluorescence of chlorophyll, which is absent from mature functional heterocyst cells) of *Nostoc* 7120 substrains, 48 hours after nitrogen stepdown. White arrows indicate heterocyst cells. Scale equals 10 µm.

Nitrogen fixation is performed in the heterocyst cells, so the observed reduction in nitrogen fixation can result from multiple factors: (i) a reduction in heterocyst frequency; (ii) heterocyst malfunction, caused by various reasons, such as impaired heterocyst cell envelope that disrupts the heterocyst microoxic environment or altered induction of genes affecting nitrogen fixation; or (iii) a combination of both. To assess whether resistance resulted from reduced heterocyst frequency, the heterocyst differentiation in the resistant substrains was compared to that of the susceptible control. A significant reduction was identified in heterocyst frequency for all of the examined *Nostoc* 7120 (Fig. 3B, S3, Table 1). However, the specific heterocyst frequency in the different resistant substrains varied. For example, while substrain RD2 completely lost the ability to induce heterocysts, substrain RE1 had a much smaller reduction (27%) in heterocyst frequency (Fig. 3B), relative to the susceptible paired controls. While these results suggest that the loss of nitrogen fixation was due to the reduction in heterocysts frequency, the only partial reduction observed in most substrains suggests that other factors likely contribute to the observed nearly complete loss of N_2_ fixation activity. As mentioned above, reduced nitrogen fixation can happen due to malfunction of the heterocysts. Changes to the heterocyst that lead to such malfunction can sometimes be visualized by microscopy, but in other cases cannot be visualized (Fig. 3D). To understand whether the loss of nitrogenase activity is a consequence of low expression of the nitrogenase enzyme, we analyzed the expression of *nifH* (which encodes for nitrogenase reductase), at 48 hours post nitrogen stepdown. The *nifH* expression was significantly decreased in all of the analyzed substrains (13/13) and varied between 3-45% of that of the susceptible control (Fig. 3C, S3B, Table 1). These results suggest that the loss of nitrogenase activity in the tested mutants is a result of different mechanisms: loss of heterocyst induction, or a combination of reduced heterocyst appearance and heterocyst malfunctioning.

### Mutations in the resistant substrains

The previous analyses show that resistance to phages is coupled with a lower ability to fix atmospheric nitrogen and/or to survive under nitrogen starvation. Furthermore, we observed a variety of phenotypes in the resistant substrains, suggesting that different mechanisms are involved in this process. To better characterize these mechanisms, we used whole genome sequencing to identify the pleiotropic mutations present in the phage resistant *Nostoc* 7120 substrains.

We sequenced the genomes of 58 *Nostoc* 7120 substrains resistant to phage A-4L and/or AN-15, as well as resequenced five genomes of their susceptible ancestors. We identified 123 mutations common to both resistant and susceptible substrains (Table S2), likely acquired prior to the initiation of this experiment and unlikely to be related to phage resistance (hereafter control mutations). We also identified 50 additional mutations that were specific to the resistant substrains (hereafter resistance related mutations; Table S1). Eleven of the resistance-related mutations repeated in different resistant substrains originated from the same and/or from different ancestral substrains. Most of the resistance related mutations (33/50) were single nucleotide point mutations (SNPs), while the others were insertions or deletions (Table S1). Most of these mutations were found in coding regions (42/50) and were non-synonymous (34/50; Table S1). Most of the mutations were found in or upstream to non-core genes (32/50), which corresponded to the proportion of non-core genes in the genome of *Nostoc* 7120 (Fisher’s exact test, *p*=0.752; Fig. 4). One of the mutations that were identified within an intergenic region, was located in a spacer region within a CRISPR array (RE8; Table S1). Spacers present in CRISPR arrays that are identical or nearly identical to the corresponding protospacers within phage genomes, take part in the host’s defense against phages, mediated by the CRISPR-cas system [20]. However, the mutant spacer within the RE8 array demonstrated no sequence similarity to the genomic sequence of the phage used for selection (A-4L). Therefore, it is unlikely that this mutant spacer confers resistance against A-4L. RE8 harbored five additional mutations (Table S1), one of which is probably responsible for the phage resistance.

**Figure 4.**
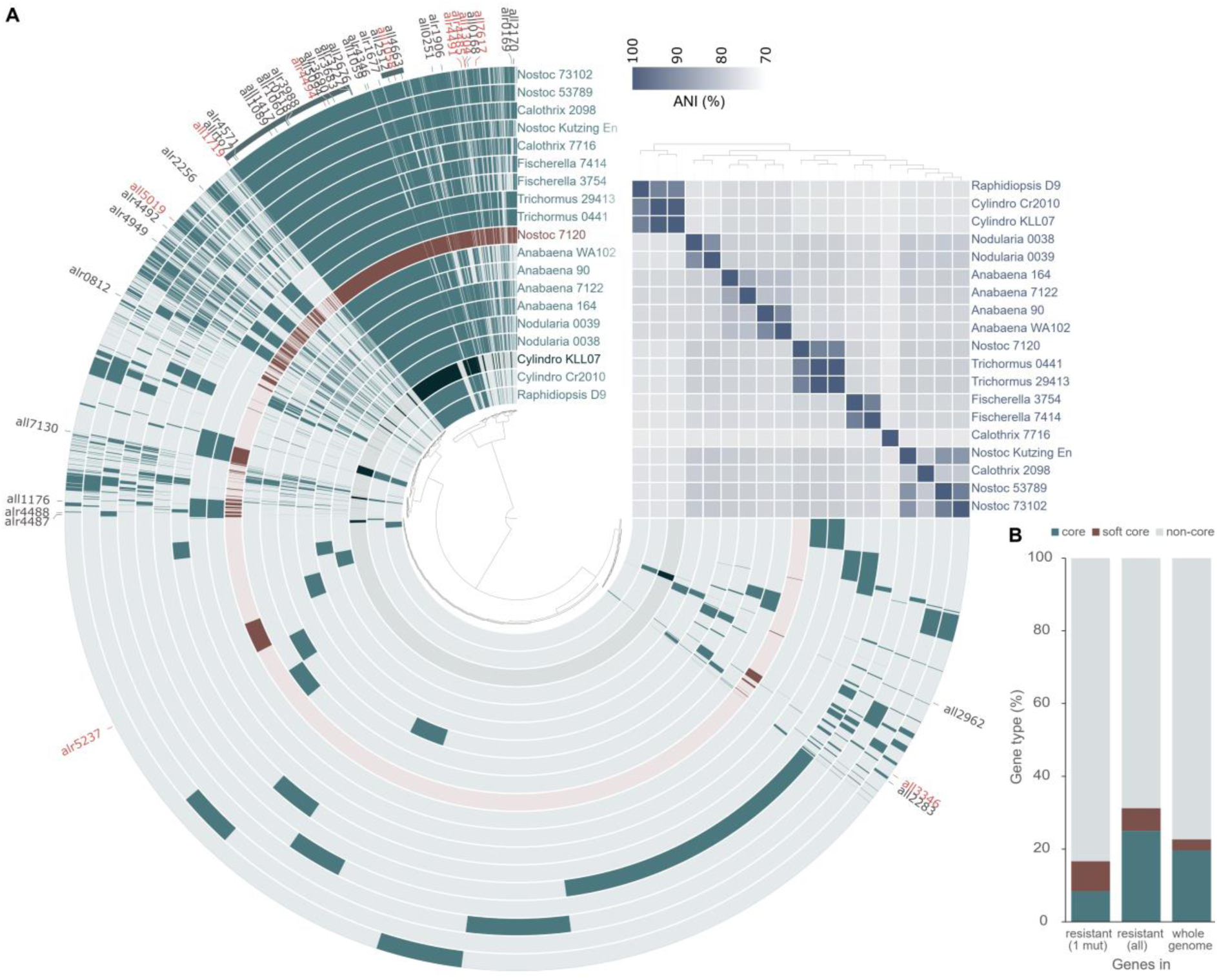
Core and accessory genome of *Nostoc* 7120. **A**. Pangenome analysis and average nucleotide identity (ANI) for 19 *cyanobacteria* strains belonging to the *Nostocales* order (full names and accession numbers are listed in Table S5). The inner tree illustrates the relationships between 23,526 gene clusters identified across these cyanobacteria genomes. Each layer of the circle phylogram represents a different organism, and the presence of a gene cluster for each organism is depicted by the darkness of the respective layer. The layers representing *Nostoc* 7120 and *C. raciborskii* are highlighted in brown and dark green, respectively. The outermost partial layer in dark green signifies gene clusters found in at least 18 out of the 19 analyzed genomes, representing the “soft core” genome. Within this collection of gene clusters, there are two distinct subgroups. The larger subgroup, situated on the left, comprises gene clusters that are universally present in all 19 genomes, thus categorizing them as "core" gene clusters. The locations of 42 gene clusters harboring genes that were found to be affected by mutations associated with resistance are indicated outside the circle phylogram. These genes are color-coded to distinguish between those that underwent mutation in substrains with a single mutation, which are highlighted in orange, and the remaining genes, depicted in grey. The ANI percentages are presented as a heatmap (color scale: top right corner). The hierarchical arrangement of the tree above the heatmap is structured based on the ANI percentage values. **B.** The distribution of core genes, soft core genes, and non-core genes within the mutants exhibiting a single mutation, across all the mutants, and throughout the entire genome of *Nostoc* 7120. Statistical analysis revealed no significant difference between the observed gene distribution in the resistant mutants and the predicted values across the entire genome.

A total of 42 genes were found to be affected by mutations associated with resistance. Among these genes, 18 had cell surface-related functions, including various functions such as transporters and enzymes crucial for cell surface biosynthesis (Table S3). Ten of the mutated genes were classified as regulatory genes based on their annotations and/or their observed phenotypes, such as the “Mch” phenotype characterized by the presence of two or more adjacent heterocysts. One of the regulatory genes (alr4494) was a glycosyl transferase involved in cell surface biogenesis, while strains with a single mutation in this gene (RB1, RB8, RB9, and RB20) exhibited the Mch phenotype (Table 1), suggesting this gene has a regulatory role. Among the regulatory genes, all2512 was identified as encoding PatB, which is an important regulator in heterocysts maturation [48]. The remaining mutant genes had other known (10) or unknown functions (4; Table S3).

### Genes in which mutations occurred multiple times

Mutations occurring multiple times within the same gene, particularly under specific selection pressures, often signify the gene’s important role in responding to the applied selection. We identified several instances of recurrent mutations in seven distinct genes (Fig. 5, Table S1). Three of these genes (all1304, all3346, and alr4491) exhibited a distinct mutation pattern characterized by mutations unique to the resistant strains, as well as mutations shared between resistant and control strains. In all3346 and alr4491, the mutations observed in the control strains were silent (Table S1), while the mutations specific to the resistant strains were located upstream of the gene or represented non-synonymous changes. This may suggest that the mutations found in the control strains did not significantly impact the cellular phenotype. Moreover, the gene all1304, responsible for encoding the bicarbonate transporter BicA, displayed an additional mutational profile that paralleled the patterns observed in the previously discussed genes. In this gene, two missense mutations were identified in the control strains, while all three resistance-associated mutations observed within all1304 were nonsense mutations (indels). These findings further emphasize the critical role of mutation characteristics in the context of resistance.

**Figure 5.**
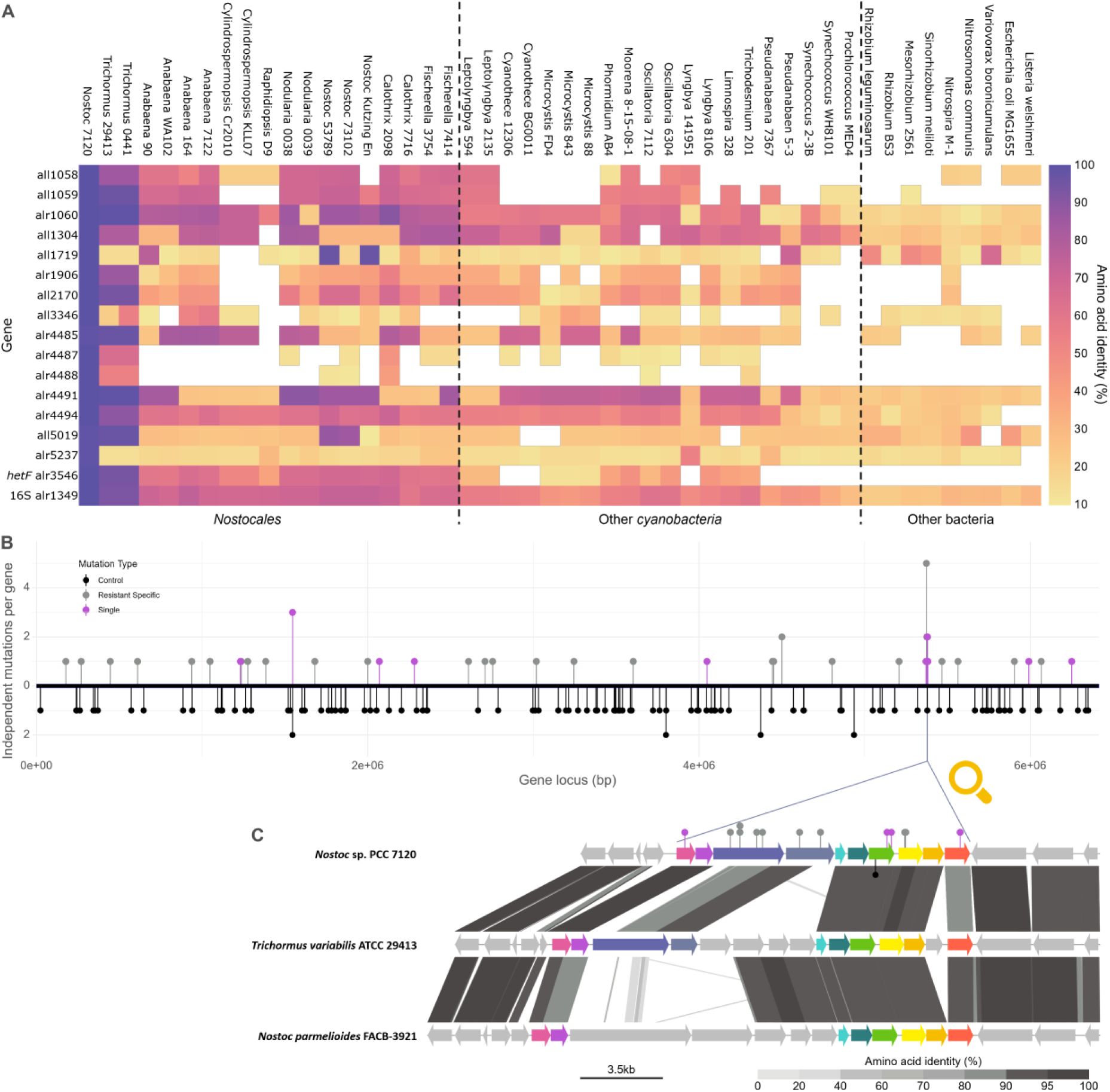
Genome evolution of *Nostoc* 7120. **A**. Heatmap of orthologs of genes involved in resistance to phages and/or in nitrogen fixation in *Nostoc* 7120 presented as average % identity of amino acid sequences. The heatmap includes protein products of *hetF* (alr3546) and 16S rRNA (uracil(1498)-N(3))-methyltransferase (alr1349) as references. The hierarchy of bacteria used for the analysis is ordered using maximum likelihood phylogenetic tree of alr1349. **B**. A schematic representation of mutations along the chromosome of *Nostoc* 7120, in the susceptible wild types (black markers), in resistant mutants with a single mutation (purple markers), and in resistant mutants with multiple mutations (grey markers). Each position along the horizontal axis corresponds to a different gene within the genome of *Nostoc* 7120, and the vertical axis provides information about the number of mutations that independently occurred in each gene. **C**. Genomic neighborhood of a cell surface-related gene cluster (alr4485-alr4494), in which mutations were found in 35 of the resistant strains. The genomic composition (at the amino acid level) within this gene cluster and its flanking regions were compared to the corresponding regions in close relatives of *Nostoc* 7120. Colored genes (other than grey) represent genes within the cluster in *Nostoc* 7120 and their homologs. Each color corresponds to a different COG number. Colored markers represent mutant loci within this cluster. Color code of the markers as in (B).

Four of the genes, in which multiple mutations were identified (alr4487, alr4488, alr4491, and alr4494) were found within a single gene cluster (Fig. 5), which is enriched with genes associated with cell surface functions. Further analysis and discussion of this gene cluster are provided below. While the presence of multiple mutations within a single gene emphasize the importance of that gene in adapting to the selective environment, the fact that the majority of the affected genes exhibited only a single mutation (36/43), suggests that our current understanding of the genetic basis underlying resistance in *Nostoc* 7120 against phages A-4L and AN-15 remains incomplete.

### Genes involved in resistance to A-4L and AN-15 and in nitrogen fixation

The number of mutations present in each resistant substrain varied. Fifteen of the 58 resistant substrains had a single mutation in their genome, 26 substrains had two mutations, and nine substrains had three or more mutations. The remaining eight genomes, in which 1-3 mutations were identified, had low coverage, and thus we could not exclude the possibility that additional mutations may be present in these genomes. In substrains carrying multiple mutations, it is impossible to identify which is the mutation that is responsible for the phage resistance and/or the reduction in nitrogen fixation without further genetic manipulation. Therefore, we focused our discussion of genome evolution on the 15 substrains carrying a single mutation (Table 1, S1, S3).

Only one of these 15 substrains (RE4) carried a mutation in a locus that was previously connected to either nitrogen fixation or to heterocyst induction. This mutation was found in a gene encoding for CusA/CzcA family heavy metal efflux RND transporter (all7617) that was up regulated during nitrogen starvation [49]. The other mutant genes, which have not previously been linked to nitrogen fixation and/or heterocyst induction, were classified into two groups: cell surface-related and/or regulatory genes (Table S3).

Twelve of these 15 substrains carried mutations in cell surface-related genes. Most of these genes encode for proteins involved in cell wall biogenesis or modification, transport of cell surface molecules or proteins embedded in the outer membrane (Tables 1, S1, S3). Molecules located on the cell surface can serve as receptors of various phages [43, 50, 51], so these mutations may alter the structure of phage receptors and thereby prevent or reduce phage adsorption [43, 50, 52]. Additionally, mutations in cell surface genes may alter the heterocyst cell envelope, increasing the permeability of these cells and interfering with their micro-oxic environment, thus preventing nitrogen fixation by inactivating the nitrogenase complex and/or by down regulating the expression of *nifH* [6, 8, 9].

The other substrains had mutations in or upstream to genes that seem to have unknown regulatory functions. For example, substrain RD1 has a single mutation in all1719, which encodes for a DNA-binding transcriptional regulator (Table 1, S1, S3). This mutation is associated with a reduced heterocyst induction (27%; Table 1, Fig. 3B) and nearly no nitrogenase gene expression (4%) and activity (3%). This suggests that this gene takes part in the regulation of heterocyst differentiation as well as nitrogen fixation in the induced heterocyst cells. However, this substrain continues to grow under nitrogen starvation, which is quite striking due to its negligible nitrogenase activity. Two other mutations were identified in two substrains that both form heterocysts with a deformed morphology, which sometimes appeared in groups of 2-3 adjacent cells in different stage of differentiation (Fig. 3D, S3). This appearance resembles the multiple contiguous heterocyst (Mch) phenotype [53–56] that was observed previously. One of these substrains (RE1) had a mutation in all5019, a gene that encodes for 3-phosphoshikimate 1-carboxyvinyltransferase, which is involved in aromatic amino acids biosynthesis. The other substrain (RE3) has a mutation located upstream to all3346, which encodes for a Repeats-in-toxin (RTX) toxin-related Ca^2+-^binding protein. RTX proteins are a family of proteins secreted via a type I secretion system. The Mch phenotype is known to result from the abnormal expression of various regulatory genes, such as overexpression of the genes *hetF, hetR*, *hetP*, and *hetZ* [56–59], or by inactivation of one of the genes *patS* and *hetN* [54, 60]. Although the exact regulatory pathway in which these genes take part of is yet unknown, the Mch phenotype caused by mutations in all5019 and upstream of all3346 suggest that they are potential regulators of heterocyst differentiation. A similar phenotype was observed in three substrains (RB1, RB8 and RB20) carrying a single mutation in alr4494, which encodes for a glycosyltransferase involved in cell wall biosynthesis (Table 1). This suggests that this gene might function both as cell surface and regulatory gene.

In addition to these regulatory genes identified in resistant substrains with a single mutation, we also identified another potential regulatory gene in substrain RD2, which carries two mutations (Table 1, S1, S3). While the presence of a second mutation in the genome of RD2 complicates the analyses, our data suggest that the loss of the ability to fix nitrogen in this strain is conferred by the mutation in all2170, which encodes for a “Caspase HetF Associated with the Tetratricopeptide repeats” (CHAT) domain-containing protein. HetF (encoded by alr3546) is an essential regulator of heterocysts differentiation in *Nostoc punctiforme* and in *Nostoc* 7120 [55, 56, 61]. Both HetF and all2170 belong to the same protein family (pfam12770). The phenotype of this strain is very similar to that of *ΔhetF* mutants as well as mutants with single amino acid substitutions in *hetF* [55, 56, 62]: (i) lack of heterocyst induction under nitrogen starvation (Fig. 3A and 3D); (ii) significantly larger cells (Fig. 1A, S1A); (iii) aberrant cell morphology (Fig. 3D); and (iv) highly granulated cells. Therefore, we propose that all2170 and HetF are involved in the same regulatory pathway that controls heterocysts differentiation.

We note that the two substrains carrying a single mutation in a regulatory gene that were tested for the mode of phage resistance (RD1 and RE1) both displayed a significant reduction in phage adsorption (Table 1). This result suggests that resistance to phages in these substrains is due to cell surface modifications downstream of the mutant regulatory genes.

### Phage selection increase the heterogeneity of genes that affect heterocyst formation and functioning

Nitrogen fixation in diazotrophic bloom-forming *cyanobacteria* is a highly advantageous trait that enables bloom formation. Therefore, it may be expected that genes that are essential for nitrogen fixation should be conserved in diazotrophic *cyanobacteria*. However, phylogenetic analyses of the mutant genes described above and their homologs in 19 heterocystous *cyanobacteria* of the *Nostocales* order as well as 29 substrains of other cyanobacterial and bacterial orders suggest that this is not necessarily the case (Fig. S4). While *hetF*, a well-known regulatory gene essential to heterocyst formation and thus to nitrogen fixation, has homologs with amino acid identity greater than 30% to the *Nostoc* 7120 gene, in all other 18 analyzed *Nostocales* strains (Fig. 5A), the genes identified in this study have homologs in fewer strains (ranging from 0-18) and these homologs display lower amino acid identity (Fig 5A). For example, only two of these genes, cluster with all (alr4494) or the vast majority (17/18, all1058) of the other *Nostocales* strains and belong to the *Nostocales* core (or soft core) genome (Fig. 4A). However, other genes, such as all1719 (Fig. S4E), cluster with a few orthologs in other *Nostocales* strains as well as heterotrophic bacteria, suggesting they were acquired by lateral gene transfer. In many genes that have homologs in other *Nostocales* strains, the clustering pattern suggests that they went through multiple lateral gene transfer events between various *cyanobacteria* strains (Fig. S4). These results suggest that genes that are essential for heterocyst induction and functionality are not necessarily core genes, and their phylogeny is affected by lateral gene transfer events. It is possible that in other *Nostocales* strains, the function of these genes is fulfilled by other genes.

Further evidence of the phage selection’s impact on the evolution of genes essential for nitrogen fixation was the identification of multiple mutations within 6 out of 10 genes located in a long gene cluster that is enriched in cell surface-related genes (alr4485-alr4494; Fig. 5C). Thirty-five of the resistant *Nostoc* 7120 substrains carried a mutation in this cluster, and in seven of the substrains this was the only mutation identified (Table S1). Most of the mutant genes in this cluster are *Nostocales* non-core genes (Fig. 4A, 5A, S4, Table S3). Moreover, a comparison with this genomic region in other *Nostocales* strains, closely related to *Nostoc* 7120, suggest that this region experienced multiple lateral gene transfer events (Fig 5C). Therefore, our results suggest that cyanophages select for the diversity of heterocyst related genes, which is similar to previous studies proposing that cyanophages select for the diversity of genomic islands in marine *cyanobacteria* [25, 63] and in other bacteria [63].

### Similar tradeoff between phage resistance and nitrogen fixation in the bloom-forming *cyanobacteria C. raciborskii*

To investigate whether the tradeoff between resistance to phages and nitrogen fixation extends to other heterocystous *cyanobacteria*, we examined *C. raciborskii*, an invasive member of the *Nostocales* with broad geographical distribution and known to form harmful blooms [14, 64–67]. For this, we used a *C. raciborskii* strain that was previously isolated from Lake Kinneret, Israel (where *C. raciborskii* forms annual blooms) [68] and also isolated three novel phage strains from a summer bloom in Lake Kinneret, called Cr-LKS4, Cr-LKS5, and Cr-LKS6 (see supplementary data). We then isolated 8 substrains of *C. raciborskii* resistant to these three phages and assessed their growth under nitrogen replete and nitrogen limitation conditions, which was compared to that of their parental strains.

When grown in a nitrogen rich medium, none of the resistant substrains (0/8) showed reduced growth (Fig. 6A, S2). However, in nitrogen depletion conditions, all of the resistant substrains (8/8) collapsed, while the controls survived (Fig. 6B; S2). Moreover, a significant reduction was identified in heterocyst frequency for all of the examined *C. raciborskii* resistant substrains (5/5; Fig. 6C). Working with *C. raciborskii* and its phages was technically challenging, and thus additional physiological characterization of this tradeoff was not carried out. We further sequenced the genomes of these eight resistant substrains and their paired controls, but were unable to identify mutations specifically linked to resistance, likely due to the high activity of transposable elements in these strains. These transposable elements may disrupt genes that are essential both for phage infection and nitrogen fixation. However, tracking the movement of these elements can be challenging using standard methods. Therefore, elucidating the mechanisms behind the tradeoff between resistance to phages and nitrogen fixation in *C. raciborskii* will need to be pursued in future studies. Nonetheless, these data demonstrate that the tradeoff between phage resistance and nitrogen fixation is observed across multiple heterocystous *cyanobacteria*, including not only the model *Nostoc* 7120 but also the bloom-forming species *C. raciborskii*.

**Figure 6.**
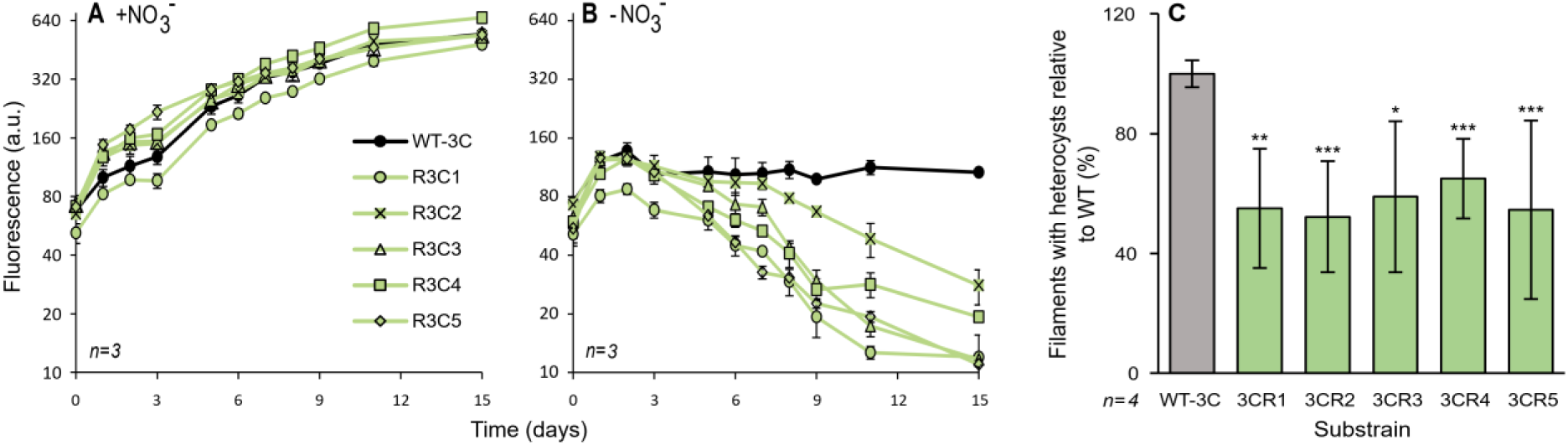
Cost of resistance in *C. raciborskii*. Growth dynamics of resistnat substrains and their susceptible controls in nitrogen replete (**A**) and nitrogen deplete conditions (**B**). The difference in growth over time for all substrains grown in nitrogen starvation conditions was significaltly lower than that of the susceptible wild type (*p*<0.01 for R3C2 and R3C4, and *p*<0.001 for all the rest). However, when grown in nitrogen rich medium, the resistant substrains had no significant reduction in growth with respect to the susceptible wild type. All resistant substrains had a significantly greater cost under nitrogen starvation than in nitrogen replete medium (*p*<0.05 for R3C2 and R3C4, *p*<0.01 for R23C1 and R3C3, and *p*<0.001 for R3C5). Data shown are average +/- standard deviation of three biological replicates. Relative cell density is estimated by chlorophyll *a* autofluorescence. a.u: arbitrary units. **(C)** Percentage of filaments with heterocysts of the resistant substrains, relative to the susceptible wild type (grey), 6 days after nitrogen stepdown. Data shown are average and standard deviation of four biological replicates. * *p*< 0.05, ** *p*< 0.01, *** *p*< 0.001.

### Conclusions

In this study, we examined host:phage interactions in two species of diazotrophic *cyanobacteria*: *Nostoc* sp. and *C. raciborskii*. Our results show that *cyanobacteria* can evolve phage resistance that is associated with a fitness tradeoff by which resistant substrains have reduced ability to fix nitrogen and/or to survive nitrogen starvation. We identify several mutations associated with this tradeoff, including in genes related to cell surface and genes regulating heterocysts development and function. Furthermore, we provide evidence that most of these genes are accessory genes whose evolution is likely impacted by phage infections.

While nitrogen (N_2_) is the most abundant element in our atmosphere, it is not available to most organisms on earth [1–3]. The ability to fix atmospheric nitrogen is highly beneficial to diazotrophic *cyanobacteria* species during seasons of low nitrogen availability in their environment. This ability gives these species an adaptive advantage over other phytoplankton species and enables them to form blooms worldwide [10]. Diazotrophic *cyanobacteria* coexist with the phages that infect them and can quite rapidly evolve resistance to these phages by acquiring spontaneous mutations that alter the cell surface of the host and reduce the rate of adsorption of the phage [25–30]. However, in the heterocystous *cyanobacteria* studied in this work, this resistance comes with various phenotypes, such as reduction in heterocyst induction, non-functional heterocyst cells, or a complete loss of heterocyst induction that eventually lead to reduced or even complete loss of the ability to fix nitrogen and thus to survive in nitrogen starvation conditions.

Our genomic analyses suggest that this pleiotropy is a result of mutations that cause, directly or indirectly, changes to the structure of the cell surface of the *cyanobacteria*. These changes probably lead to the reduced phage adsorption that was observed, and thus to phage resistance. Most of the mutations were found in cell surface-related genes, such as transporters of sugars and other enzymes that take part in cell wall or membrane formation and modification. The structure integrity of the cell wall of the heterocyst cell is crucial for the microoxic environment of the heterocyst, which is required for nitrogenase expression and function. Moreover, cell surface aberrations can lead to fractionation of the filaments next to the heterocysts, which may cause heterocyst loss [69]. Other mutations were found in regulatory genes that caused the complete loss or irregular induction of heterocyst cells, as well as other unknown changes that resulted in lack of nitrogen fixation. The fact that RE1 that had a single mutation in a regulatory gene showed impaired adsorption, suggested that one of the downstream effects of these mutations is cell surface modification.

The ability to fix nitrogen is one of the central traits that enables heterocystous *cyanobacteria* to bloom under nitrogen starvation conditions. Therefore, the tradeoff between resistance to phages and nitrogen fixation may affect *cyanobacteria* population dynamics during blooms. The fact that bloom-forming heterocystous *cyanobacteria* coexist with their phages in nitrogen starvation conditions, suggests that the *cyanobacteria* may have solutions to this paradox (e.g. other resistance conferring mutations with no tradeoff with nitrogen fixation or compensatory mutations) that enable resistance while maintaining a functional nitrogen fixation system. However, this may lead to selective pressure formed by phages on genes essential for nitrogen fixation. Indeed, the phylogeny of the mutant genes suggests that they went through multiple lateral gene transfer events.

Our findings also raise important considerations for the role of phages in managing harmful cyanobacterial blooms. While there are several limitations linked to the application of phages to address harmful blooms in large aquatic environments, such as the potential release of toxins during bacterial lysis, a better understanding of bloom dynamics, drivers, and of the factors promoting blooms decline is very important for lake management. Our findings provide valuable insights into the factors influencing the dynamics of harmful blooms under nitrogen-starved conditions. Moreover, we suggest that phage selection could serve as a valuable approach to discover genes previously unknown to be associated with the regulation and development of heterocysts. Further research focusing on the molecular mechanisms through which such genes participate in the nitrogen-fixing ability of heterocystous *cyanobacteria* will enhance our understanding of this complex process, which plays a pivotal role in blooms formation under nitrogen starvation.

## Materials and Methods

### Culturing conditions

*Cyanobacteria* were cultured in BG11 medium [70] with 2.5-3 µmol/m^2^s photons flux, under daily cycle of 10-hour dark and 14-hour light, at 24 ^°^C. For nitrogen starvation conditions, cultures were grown in BG11_0_, which lacks the main nitrogen source in BG11, NO_3-_. The transfer of cultures from nitrogen rich to nitrogen deprived medium was performed by three cycles of centrifugation (1300 g for 5 min), followed by pellet resuspension with BG11_0_ or BG11 for control. The growth of the cultures and infection dynamics in liquid medium was monitored by chlorophyll *a* autofluorescence, which served us as a proxy for the number of *cyanobacteria* cells in the culture. The measurements were performed using a Synergy 2 plate reader (BioTek) with excitation and emission wavelengths of 440nm and 680nm respectively.

*Cyanobacteria* single colony isolation was performed by pour plating, following Avrani et al. (2011) with minor modifications. In short, *cyanobacteria* cultures in logarithmic phase were ten-fold serially diluted and poured with BG11 medium and low melting point agarose at a final concentration of 0.28% into 90 mm Petri dishes.

### Cyanobacteria strains

Two cyanobacterial strains of the order *Nostocales* served as models for filamentous bloom-forming *cyanobacteria*. The first is *C. raciborskii* st. KLL07, which is nontoxic and was isolated from Lake Kinneret [68]. The second strain is the model strain *Nostoc* sp. PCC 7120.

### Cyanophage strains and isolation

To select for resistance in *Nostoc* 7120 we used the phage A-4L that was isolated in 1972, Leningrad, USSR [71] and the phage AN-15 that was isolated in 1981 in south-central Michigan [39]. These phages were obtained from the ATCC culture collection. The lysates that used to infect *Nostoc* 7120 included A-4L, AN-15, or a mixture of both phages. To select for resistance in *C. raciborskii* we isolated three new phages (Cr-LKS4, Cr-LKS5, and Cr-LKS6) from station A in Lake Kinneret (32°82.146′N 35°35.191′E), which is the same location from which the host strain was isolated *7* years earlier. These phages were isolated from the same water sample and in a similar method as Cylindrospermopsis phage Cr-LKS3 [72]. In short, water was sampled at a depth of 5 m on September 5, 2017 at the IOLR monitoring station at the center of Lake Kinneret. Isolation was performed by four sequential plaque assays with *C. raciborskii* st. KLL07 as the host. Plaque assays were performed by pour plating 2.8 ml of host population (*C. raciborskii*) in mid-log phase with 200 µl of ten-fold serially diluted phage containing lysate, together with BG11 low melting point agarose at a final concentration of 0.28%. The isolated phage was used to inoculate the host in liquid medium, and the resulting lysate was filtered through a 0.22 μm filter.

### Isolation of *cyanobacteria* substrains resistant to phages

In order to isolate resistant substrains carrying a minimal number of mutations differentiating them from their paired susceptible control, phage selection was generated on isogenic populations that originated from single colonies. Isolation of the resistant substrains was performed using two selection methods: selection in liquid medium and selection in semi-solid medium (Table S1, S4).

Selection in liquid was achieved by infecting the susceptible WTs in mid-log phase in a 96 well plate with the selecting phage or without the phage for control. The infection dynamics was monitored by a fluorescence plate reader. While the uninfected controls continued to grow, the infected populations declined due to phage infection. The collapse of the infected cultures was followed by a population recovery. A sample of the evolved recovered population was ten-fold serially diluted and pour plated to form single colonies that subsequently were transferred into a liquid medium for further analysis.

For selection in semi-solid medium, we followed the procedure used by Avrani et al, (2011) with minor modifications. A susceptible culture originated from a single colony (susceptible WT) in mid-log phase, was serially diluted. The dilutions were pour-plated with the selecting phage for resistant substrain isolation, and with no phage for control. The isolated colonies of the resistant substrains (selected by the phage) were transferred to a liquid medium for further analysis.

The resistance of the isolated substrains was validated by reinfection by the initial selecting phage using susceptibility tests as follows: for each culture, three wells with 180 µl of *cyanobacteria* culture in mid log phase were inoculated with 20 µl of the selecting phage. Three uninfected wells (180 µl of *cyanobacteria* culture added with 20 µl of BG11) served us as negative controls. Similar experiment with the susceptible paired control of each strain served as a positive control for phage infection. The growth of the cultures was monitored for a week after the susceptible cultures collapsed.

### Cell sizes measurements

Cultures in mid-log stage were sampled and imaged using an inverted bright-field (BF) microscope, Nikon Eclipse Ti2-E. The size measurements were carried out using NIS-Elements AR Analysis 5.11.01 software. Cells were sampled from at least two filaments and were measured parallel and perpendicular to the growth axis (width and length respectively). We chose 11 substrains belonging to three different lineages (B, D and E). In each lineage, strains that showed different colony phenotypes (color, aggregation) were chosen. Where there was no intra-lineage variation (as in the lineage B), the substrains were chosen randomly.

### Adsorption assays

Prior to the adsorption assays the cultures were isolated from the selecting phage first by three cycles of centrifugation (1300 g for 5 min), followed by pellet resuspension with BG11, and by serial dilutions of the pellet that were pour plated to form single colonies. These colonies were subsequently transferred into a liquid medium for further analysis. We then verified that these cultures had no phage in them, using spot assays following Avrani et al. (2011). Spot assays were performed using a mixture of BG11 and low melting point agarose in a final concentration of 0.28%. The mixture was pour plated with a susceptible host population (*Nostoc* 7120) into 90 mm Petri dishes to form lawn. Then, 10 µl of the examined culture was spotted on the host lawn to detect phage presence. All strains that were examined (RD1, RD2, RD3, RE1, RE2, and RE4) had phages in the *cyanobacteria* culture. Strains that could be isolated from the selecting phage were further analyzed. Analyses other than adsorption assays were performed using cultures that may contain phages, except for RE4. Analyses with the clean RE4 culture showed similar trend to the other analyzed stains. The *cyanobacteria* cultures were inoculated with phage in a multiplicity of infection (MOI) of 0.08 ≤ MOI < 1 with host cell concentration of 6 ∗ 10^7^ cells/mL. The cultures were sampled right after inoculation (t_0_) and a few hours later (t_max_), depending on the susceptible WT the resistant substrains have evolved from (t_max_ =2 hours and t_max_ =3 hours for the isolates of WT-D and WT-E respectively). In each time point, the samples were filtered through a 0.22 µm filter. This filtrate was used to estimate the free phage fraction (phages not associated with the host) at each time point, by plaque assays as above (Cyanophage strains and isolation section).

### Cost of resistance in growth

Cost of resistance in growth was assessed with or without combined nitrogen (incubated in BG11 or BG11_0_ respectively). We followed the growth of 17 resistant strains that evolved from 5 susceptible wild type strains, and preferably varied in their colony phenotype within each group with common ancestry. The growth of the resistant substrains was compared to the growth of their susceptible paired control substrains, in nitrogen replete and deplete conditions. Two-way repeated measure ANOVA of three time-points for the normalized values (the fluorescence values were divided by the initial value, t=0) was used to determine whether the growth of the resistant substrains was significantly slower than that of their paired control. See statistical analyses for details.

### Assessment of heterocyst induction

Heterocyst cells are induced during nitrogen starvation. To induce heterocysts, the cultures were transferred from BG11 to BG11_0_ medium as described above. Each experiment included substrains resistant to phage and their paired control. The heterocyst differentiation process has been tracked by bright field and fluorescence (excitation: 480 nm, emission: 700 nm) microscopy, using a Nikon Eclipse Ti2-E inverted microscope. For each biological replicate, 50 filaments were sampled randomly at t_0_ and on the t_end_ timepoint post nitrogen depletion (t_end_ values: 48 hours for *Nostoc* 7120 and 144 hours for *C. raciborskii*). For one strain, and one control strain we used less filaments in some of the biological replicates (37 and 26 in two out of six replicates for RD3 and 28 in one out of 6 replicates for its susceptible ancestor). The percentage of filaments carrying morphologically mature heterocysts at t_end_ were calculated.

### Assessment of nitrogenase activity

Nitrogen-fixation rates were measured as described previously [12] by assessing nitrogenase enzyme activity using the acetylene reduction assay. We analyzed a subset of four substrains, belonging to two lineages (D and E) derived from the substrains analyzed in the previous sections. The results of all four strains were nearly identical, and because this assay is highly time consuming, we did not further analyze additional strains. Culture samples of 5ml were placed in 28-mL serum bottles, sealed with a rubber septum and reinforced with an aluminum closure. The bottles were flushed for 5 min with argon followed by the addition of C_2_H_2_ (20% of head space volume), and incubated for 48 h under controlled light and temperature conditions. For each sample set, an additional bottle treated as above but covered with aluminum foil was used as a dark control. After incubation, samples were analyzed immediately for C_2_H_4_ accumulated in the sample by injection of 1 mL of the headspace gas to a GC-FID (Shi-madzu) using Durapak phenyl isocyanate on 80/100 Porasil C in a 60 9 1/8′′ column (Supelco), calibrated with ethylene (100 ppm standard, Supelco, Cat No: 22572; Sigma–Aldrich, Rehovot, Israel). Chlorophyll concentrations in all samples were determined by fluorometry after 90% acetone extraction using the method of Holm-Hansen et al. [73]. The values obtained in the dark control were subtracted from each measurement and the nitrogenase activity rates were normalized to the chlorophyll content in each sample, and averaged over three technical replicates for each biological replicate.

### Gene expression of *nifH*

To estimate the differences in gene expression of *nifH* in the resistant substrains of *Nostoc* 7120 relative to their susceptible WTs, RNA was extracted 48 hours after nitrogen stepdown for RT-qPCR. The RNA extraction was carried out using the zymo Quick-RNA MiniPrep Plus Kit (ZR-R1057) and the ZR BashingBead Lysis with an additional step of DNAse I (TURBO DNA-free AM1907). The RNA was quantified using a Qubit 2.0 fluorometer (invitogen) and was converted into cDNA using the High-Capacity cDNA RT kit (AB-4374966). Following the procedure in [74], each qPCR reaction contained 1X LightCycler 480 SYBR Green I Master mix (Roche), 500 nM desalted primers for *nifH* and *rnpB* [75] and 2 μL of cDNA template (rang between 0.5 to 1ng) in a total reaction volume of 10µl of 10 mM Tris·HCl [pH 8]. Reactions were carried out on a QuantStudio Real-Time PCR system (Applied Biosystem). The cycling program began with a denaturation step of 95 °C for 10 min, followed by 40 cycles of amplification. Each cycle consisted of denaturation at 95 °C for 15 s, annealing and elongation at 60 °C for 1 min. Afterwards, the fluorescence values of each plate were recorded (6-Carboxyfluorescein [6-FAM]; excitation/emission: 465/510 nm). The point at which the fluorescence of a sample rose above the background fluorescence was calculated using QuantStudio design and analysis software version 1.5.1.

Specificity of the amplified PCR product was verified by performing a melting curve analysis on the QuantStudio Real-Time PCR system (Applied Biosystem) instrument. Standard curves were generated using plasmids containing cloned target sequences, and used to calculate the number of gene copies in the samples. For this purpose, PCR products of *nifH* and *rnpB* were each cloned into the pGEM-T plasmid (Promega). cDNA concentrations in nanograms per microliter were measured by Qubit 2.0 fluorometer (invitogen). DNA concentrations were converted to genome copies per milliliter by entering the genome length of the pGEM plasmid into the DNA copy number and dilution calculator by Thermo Fisher Scientific Inc for determining the number of copies of a template (https://www.thermofisher.com/il/en/home/brands/thermo-scientific/molecular-biology/molecular-biology-learning-center/molecular-biology-resource-library/thermo-scientific-web-tools/dna-copy-number-calculator.html).

The number of copies of the target gene *nifH* were normalized using the number of copies of the *rnpB* gene which served as a reference gene. To achieve the relative normalized copy number of the *nifH* gene, values were divided by the normalized copy number of *nifH* in the susceptible paired control.

### Statistical analyses

To assess a significant difference between the sizes of the cells (width and length) in the resistant substrains and in their susceptible WTs, two tailed t-tests were used. To assess significant difference between the resistant substrains and their susceptible WTs in adsorption of phage particles to the host cell surface, filaments with heterocysts, nitrogenase activity, and normalized *nifH* expression, one tailed t-tests were used. All of the above was done when the data were normally distributed according to *p*>0.05 using the Kolmogorov-Smirnov or Shapiro-Wilk test. In cases where the data were not normally distributed, the non-parametric Mann-Whitney test was used instead. The latter included four cases in the width distribution (RD2, RD3, RE1, and RE4), seven cases in the length distribution (RD1, RD2, RD3, RE1, RE2, RB8, and RB20), two cases in the number of filaments with heterocysts (RD2 and R3C4), and one case when estimating the difference between the expression of *nifH* to the number of filaments with heterocysts (RE2).

#### Growth (with or without combined nitrogen)

To assess the differences in growth with respect to time between the resistant and the susceptible substrains, two-way repeated measures ANOVA tests in three time points (t=0, last day of experiment, midpoint between the two timepoints) were carried on the normalized fluorescence intensity values. The normalized fluorescence intensity was achieved by dividing the fluorescence values with the initial fluorescence value (in t=0) for each culture. The time points that were used for the analysis of the isolates of *Nostoc* 7120 are: RA2: 0, 6, 13; RB1, RB8, RB20, RD4, RE3-5: 0, 7, 14; RD3, RE4, RF2, RF22, RF25: 0, 6, 12; and RD1, RD2, RE1, RE2: 0, 4, 10. The time points used us for the analysis for the isolates of *C. raciborskii* are: R3C1-5: 0, 7, 15 and R8D1-3 0, 7, 14.

#### Growth +N vs. Growth -N

The difference in the cost in growth of the resistant substrains under nitrogen replete vs. deplete conditions was tested using two-way repeated measure ANOVA as follows: (i) the average of the normalized chlorophyll auto fluorescence values (as described above) in each analyzed timepoint was calculated in each WT strain; (ii) the corresponding values of the resistant substrains were divided by the average value of the WT for each timepoint (this was performed separately for cultures grown under nitrogen replete and deplete conditions); and (iii) two-way repeated measures ANOVA tests in the three time points was performed on the values calculated in (ii).

To assess the statistical significance of the differences between gene types (core and non-core genes) in the entire genome of *Nostoc* 7120 and in the mutated genes found in resistant substrains of *Nostoc* 7120, Fisher’s exact test was employed.

All the statistical analyses were carried out using IBM SPSS Statistics for Windows, Version 27.0. Armonk, NY: IBM Corp.

### Resequencing of *cyanobacteria* genomes

Genomic DNA of the resistant substrains and their paired controls was extracted using Presto Mini gDNA Bacteria Kit (Geneaid). Multiplexed DNA libraries were prepared using Nextera DNA Flex Library Prep Kit and following the procedure in [76]. The whole genome has been sequenced on a NovaSeq SP system (Illumina, 2X150 bp).

Adapter trimming and quality control of the data has been achieved using Trim Galore 0.4.2 (parameters: phred 33, quality 20) Resequencing and mutation calling was performed using the breseq pipeline [77] with default parameters (consensus mode with consensus frequency cutoff of 0.8 and consensus score of 10). Genomes used as reference are *Nostoc* sp. PCC 7120 (GenBank assembly accession: GCA_000009705.1) and *C. raciborskii* st. KLL07 (GenBank assembly accession: GCA_021650815.1). Identifying the mutations conferring resistance in the resistant substrains was achieved by depleting the mutations that are common to the resistant substrains and their susceptible paired controls. Identified mutations were verified by visualizing the BAM files using IGV [78]. Eight genomes had a low average coverage, which allowed the identification of 1-3 mutations in each of these genomes (local coverage in the mutation locus was 5 or higher). However, many other regions of the genome of these strains were with no coverage, and thus we could not exclude the possibility that there are additional mutations in the genome.

### Annotation of resistance related genes

Following Avrani et al. (2011) we applied several tools to annotate the resistance related genes. Annotation was relied on data from GenBank, protein family (Pfam) database [79], and by the clusters of orthologous genes (COG) database [80]. Using this information, we annotated genes as regulatory genes and/or cell surface-related genes. The annotation of a gene as cell surface-related was supported by the presence of transmembrane helices (TMH) using data from the Integrated Microbial Genomes (IMG) Data Management System [81] and by predicting localization to the membrane using PSORTb v3.0.3 [82] with minimum threshold of 7. We annotated regulatory genes also by a phenotypic appearance characterizing genes that were previously proved to be involved in regulation in *Nostoc* 7120. Moreover, we used the IMG website [81] to search for homologs in at least three other organisms and examine their genomic neighborhood which can provide further support for annotating genes as cell surface-related [83, 84]. To determine whether a mutation occurred in clustered regularly interspaced short palindromic repeats (CRISPR) sequence, we used CRISPRCasFinder [85, 86].Genes that did not fall within the categories of cell surface, regulation, or defense, yet have a different functional prediction according to the Pfam or COG databases, were classified as "other". Genes that lacked functional predictions within these databases were classified as unknown (e.g., hypothetical proteins).

### Prediction of promotor sites

Promoter sites were predicted using BPROM [87] where mutations were found in intergenic regions.

### Analysis for synonymous mutations

To study the effects of synonymous mutations, we analyzed codon usage frequency via the program *codon usage*, a part of the Sequence Manipulation Suite [88]. Furthermore, we used the minimum free energy predication of the mRNA structure using the RNAfold webserver [89, 90] to predict possible secondary mRNA structures.

### Pangenome, average nucleotide identity (ANI), and core genes analyses

The pangenome analysis of 19 *cyanobacteria* belonging to the *Nostocales* order was conducted using anvi’o v7.1 [91] and followed the pangenomics workflow [92]. Full names and accession numbers used for this analysis are provided in table S5. Prior to the analysis, gene functional annotations were imported by processing GenBank files using anvi-script-process-genbank. The pangenome calculation using anvi-pan-genome included the --enforce-hierarchical-clustering flag. Core genes and soft-core genes were identified using the filter: minimum number of genomes gene cluster occurs. The filter was set to 19 for core genes and 18 for soft-core genes. Calculation of average nucleotide identity (ANI) between genomes was carried out using the anvi-compute-genome-similarity program with a 70% cutoff.

### Phylogeny and genes heatmap

To find homologs of the examined genes of *Nostoc* 7120 for the phylogenetic analyses, we used blastp [93] search in a database comprised of 49 genomes with an e-value < 5*10^−4^. The e-value was not restricted when searching for homologs for alr1060 and all1719 due to their short sequences (150 and 105 amino acids respectively). The 49 genomes we used as a fixed database include *cyanobatecria* belonging to five different orders: *Nostocales*, *Pseudanabaenales*, *Chroococcales*, *Synechococcales*, and *Oscillatoriales* and non-cyanobacteia belonging to different groups. The list of the genomes and their GenBank or RefSeq accession numbers used by us as a database can be found in table S5. We applied an additional filter by using a cutoff of an average percent identity >10% to avoid distant sequences that have been identified by blastp as homologs using the formula 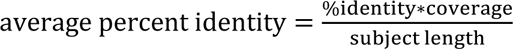 with values derived from the blastp search. Additional homologs from *Nostoc* 7120 were achieved using the same parameters.

Genes heatmap was constructed from the resulting hits using the same mentioned parameters with the Python package Plotly [94].

### Genome alignment visualization

To study the structure of the possible operon involved in resistance and nitrogen fixation in *Nostoc* 7120 and related organisms, we used the DiGAlign server (https://www.genome.jp/digalign/) with tBlastx option. The organisms used for this analysis, as well as their GenBank accessions and the interval coordinates are: *Nostoc* 7120 (GCA_000009705.1; 5356388:5389413), *Trichormus variabilis* ATCC 29413 (GCA_000204075.1; 4156514-4188547) and *Nostoc parmelioides* FACHB-3921 (GCA_014696625.1; 23892-57120 in contig8).

## Data availability

The nucleotide sequence of Cr-LKS4 has been deposited in GenBank under the accession number OQ625504.1.

## Acknowledgements

We thank Tal Dagan and her lab members, Wolfgang Hess and his lab members, Assaf Vardi and his lab members, Daniel Sher, and members of the Avrani Lab for their comments on this study, Daniel Schwartz for suggesting the title of the manuscript, and Oz Tzabari-Dar and Moti Diamant for their assistance with sample collection. We particularly thank Manuel Brenes-Álvarez for his significant advice on mutant gene function. This project was supported by the Israel Science Foundation (ISF) grant number 1386/20. Phage sampling date was determined using data from the Lake Kinneret monitoring program sponsored by the Israel Water Authority.

## Contributions

DK and SA designed the research. EKT isolated the Cr-LKS phages and performed qPCR experiments. ZF and NMR performed nitrogenase activity experiments. DK performed all other experiments. HE created genomic visualization of mutations location. SN analyzed nitrogenase activity data. DK and SA analyzed all other data. DK and SA wrote the paper with contributions from all authors.

## References

1. Berman-Frank I, Lundgren P, Falkowski P. Nitrogen fixation and photosynthetic oxygen evolution in cyanobacteria. Res Microbiol 2003; 154: 157–164.

2. Adams DG, Duggan PS. Heterocyst and akinete differentiation in cyanobacteria. New Phytol 1999; 144: 3–33.

3. Flores E, Herrero A. Nitrogen assimilation and nitrogen control in cyanobacteria. Biochem Soc Trans 2005; 33: 164–167.

4. Wolk CP, Ernst A, Elhai J. Heterocyst metabolism and development. The Molecular Biology of Cyanobacteria. 1994. Springer Netherlands, Dordrecht, pp 769–823.

5. Flores E, Herrero A. Compartmentalized function through cell differentiation in filamentous cyanobacteria. Nat Rev Microbiol 2010; 8: 39–50.

6. Nicolaisen K, Hahn A, Schleiff E. The cell wall in heterocyst formation by *Anabaena* sp. PCC 7120. J Basic Microbiol 2009; 49: 5–24.

7. Zeng X, Zhang CC. The making of a heterocyst in cyanobacteria. Annu Rev Microbiol 2022; 76: 597–618.

8. Ernst A, Black T, Cai Y, Panoff JM, Tiwari DN, Wolk CP. Synthesis of nitrogenase in mutants of the cyanobacterium *Anabaena* sp. strain PCC 7120 affected in heterocyst development or metabolism. J Bacteriol 1992; 174: 6025–6032.

9. Xu X, Khudyakov I, Wolk CP. Lipopolysaccharide dependence of cyanophage sensitivity and aerobic nitrogen fixation in *Anabaena* sp. strain PCC 7120. J Bacteriol 1997; 179: 2884–2891.

10. Gobler CJ, Burkholder JAM, Davis TW, Harke MJ, Johengen T, Stow CA, et al. The dual role of nitrogen supply in controlling the growth and toxicity of cyanobacterial blooms. Harmful Algae 2016; 54: 87–97.

11. Pollingher U, Hadas O, Yacobi YZ, Zohary T, Berman T. *Aphanizomenon ovalisporum* (Forti) in Lake Kinneret, Israel. J Plankton Res 1998; 20: 1321–1339.

12. Hadas O, Pinkas R, Malinsky-Rushansky N, Nishri A, Kaplan A, Rimmer A, et al. Appearance and establishment of diazotrophic cyanobacteria in Lake Kinneret, Israel. Freshw Biol 2012; 57: 1214–1227.

13. Bormans M. Spatial and temporal variability in cyanobacterial populations controlled by physical processes. J Plankton Res 2004; 27: 61–70.

14. Paerl HW, Otten TG. Harmful cyanobacterial blooms: causes, consequences, and controls. Microb Ecol 2013; 65: 995–1010.

15. Sigee DC, Selwyn A, Gallois P, Dean AP. Patterns of cell death in freshwater colonial cyanobacteria during the late summer bloom. Phycologia 2007; 46: 284–292.

16. Walve J, Larsson U. Blooms of Baltic Sea *Aphanizomenon* sp. (Cyanobacteria) collapse after internal phosphorus depletion. Aquat Microb Ecol 2007; 49: 57–69.

17. Bratbak G, Egge JK, Heldal M. Viral mortality of the marine alga *Emiliania huxleyi* (Haptophyceae) and termination of algal blooms. Mar Ecol Prog Ser 1993; 93: 39–48.

18. Schroeder DC, Oke J, Malin G, Wilson WH. Coccolithovirus (*Phycodnaviridae*): Characterisation of a new large dsDNA algal virus that infects *Emiliana huxleyi*. Arch Virol 2002; 147: 1685–1698.

19. Martin RM, Moniruzzaman M, Mucci NC, Willis A, Woodhouse JN, Xian Y, et al. *Cylindrospermopsis raciborskii* Virus and host: genomic characterization and ecological relevance. Environ Microbiol 2019; 21: 1942–1956.

20. Kimura S, Uehara M, Morimoto D, Yamanaka M, Sako Y, Yoshida T. Incomplete selective sweeps of *Microcystis* population detected by the leader-end crispr fragment analysis in a natural pond. Front Microbiol 2018; 9: 1–9.

21. Wang J, Bai P, Li Q, Lin Y, Huo D, Ke F, et al. Interaction between cyanophage MaMV-DC and eight *Microcystis* strains, revealed by genetic defense systems. Harmful Algae 2019; 85.

22. Chénard C, Wirth JF, Suttle CA. Viruses infecting a freshwater filamentous cyanobacterium (*Nostoc* sp.) encode a functional CRISPR array and a proteobacterial DNA polymerase B. mBio 2016; 7: 1–11.

23. Shaalan H, Cattan-Tsaushu E, Li K, Avrani S. Sequencing the genomes of LPP-1, the first isolated cyanophage, and its relative LPP-2 reveal different integration mechanisms in closely related phages. Harmful Algae 2023; 124: 102409.

24. Makarova KS, Wolf YI, Iranzo J, Shmakov SA, Alkhnbashi OS, Brouns SJJ, et al. Evolutionary classification of CRISPR–Cas systems: a burst of class 2 and derived variants. Nat Rev Microbiol 2020; 18: 67–83.

25. Avrani S, Wurtzel O, Sharon I, Sorek R, Lindell D. Genomic island variability facilitates *Prochlorococcus*–virus coexistence. Nature 2011; 474: 604–608.

26. Stoddard LI, Martiny JBH, Marston MF. Selection and characterization of cyanophage resistance in marine *Synechococcus* strains. Appl Environ Microbiol 2007; 73: 5516–5522.

27. Mole R, Meredith D, Adams DG. Growth and phage resistance of *Anabaena* sp. strain PCC 7120 in the presence of cyanophage AN-15. J Appl Phycol 1997; 9: 339–345.

28. Cowlishaw J, Mrsa M. Co-evolution of a virus-alga system. Appl Microbiol 1975; 29: 234– 239.

29. Barnet YM, Daft MJ, Stewart WDP. Cyanobacteria-cyanophage Interactions in Continuous Culture. J Appl Bacteriol 1981; 51: 541–552.

30. Yoshida T, Kamiji R, Nakamura G, Kaneko T, Sako Y. Membrane-like protein involved in phage adsorption associated with phage-sensitivity in the bloom-forming cyanobacterium *Microcystis aeruginosa*. Harmful Algae 2014; 34: 69–75.

31. Xiong Z, Wang Y, Dong Y, Zhang Q, Xu X. Cyanophage A-1(L) adsorbs to lipopolysaccharides of *Anabaena* sp. strain PCC 7120 via the tail protein lipopolysaccharide-interacting protein (ORF36). J Bacteriol 2019; 201.

32. Marston MF, Pierciey FJ, Shepard A, Gearin G, Qi J, Yandava C, et al. Rapid diversification of coevolving marine *Synechococcus* and a virus. Proc Natl Acad Sci USA 2012; 109: 4544– 4549.

33. Lennon JT, Khatana SAM, Marston MF, Martiny JBH. Is there a cost of virus resistance in marine cyanobacteria? ISME J 2007; 1: 300–312.

34. Avrani S, Schwartz DA, Lindell D. Virus-host swinging party in the oceans. Mob Genet Elements 2012; 2: 88–95.

35. Cairns J, Coloma S, Sivonen K, Hiltunen T. Evolving interactions between diazotrophic cyanobacterium and phage mediate nitrogen release and host competitive ability. R Soc Open Sci 2016; 3: 160839.

36. Padan E, Ginzburg D, Shilo M. The reproductive cycle of cyanophage LPP1-G in *Plectonema boryanum* and its dependence on photosynthetic and respiratory systems. Virology 1970; 40: 514–521.

37. Khudyakov I, Wolk CP. Evidence that the hanA gene coding for HU protein is essential for heterocyst differentiation in, and cyanophage A-4(L) sensitivity of, Anabaena sp. strain PCC 7120. J Bacteriol 1996; 178: 3572–3577.

38. Ou T, Liao XY, Gao XC, Xu XD, Zhang QY. Unraveling the genome structure of cyanobacterial podovirus A-4L with long direct terminal repeats. Virus Res 2015; 203: 4–9.

39. Hu NT, Thiel T, Giddings TH, Wolk CP. New *Anabaena* and *Nostoc* cyanophages from sewage settling ponds. Virology 1981; 114: 236–246.

40. Allen MM. Cyanobacterial cell inclusions. Annu Rev Microbiol 1984; 38: 1–25.

41. de Araujo C, Arefeen D, Tadesse Y, Long BM, Price GD, Rowlett RS, et al. Identification and characterization of a carboxysomal γ-carbonic anhydrase from the cyanobacterium *Nostoc* sp. PCC 7120. Photosynth Res 2014; 121: 135–150.

42. Flores E, Arévalo S, Burnat M. Cyanophycin and arginine metabolism in cyanobacteria. Algal Res 2019; 42: 101577.

43. Hampton HG, Watson BNJ, Fineran PC. The arms race between bacteria and their phage foes. Nature 2020; 577: 327–336.

44. Koskella B, Hernandez CA, Wheatley RM. Understanding the impacts of bacteriophage viruses: from laboratory evolution to natural ecosystems. Annu Rev Virol 2022 2022; 9: 57–78.

45. Bohannan BJM, Lenski RE. Linking genetic change to community evolution: insights from studies of bacteria and bacteriophage. Ecol Lett 2000; 3: 362–377.

46. Bennett AF, Lenski RE. An experimental test of evolutionary trade-offs during temperature adaptation. Proc Natl Acad Sci USA 2007; 104: 8649–8654.

47. Burmeister AR, Turner PE. Trading-off and trading-up in the world of bacteria–phage evolution. Curr Biol 2020; 30: R1120–R1124.

48. Liang J, Scappino L, Haselkorn R. The *patB* gene product, required for growth of the cyanobacterium *Anabaena* sp. strain PCC 7120 under nitrogen-limiting conditions, contains ferredoxin and helix-turn-helix domains. J Bacteriol 1993; 175: 1697–1704.

49. Flaherty BL, Van Nieuwerburgh F, Head SR, Golden JW. Directional RNA deep sequencing sheds new light on the transcriptional response of *Anabaena* sp. strain PCC 7120 to combined-nitrogen deprivation. BMC Genomics 2011; 12: 332.

50. Stone E, Campbell K, Grant I, McAuliffe O. Understanding and exploiting phage–host interactions. Viruses 2019; 11: 567.

51. Nobrega FL, Vlot M, de Jonge PA, Dreesens LL, Beaumont HJE, Lavigne R, et al. Targeting mechanisms of tailed bacteriophages. Nat Rev Microbiol 2018; 16: 760–773.

52. Zhang M, Qian J, Xu X, Ahmed T, Yang Y, Yan C, et al. Resistance of *Xanthomonas oryzae* pv. *oryzae* to lytic phage X2 by spontaneous mutation of lipopolysaccharide synthesis-related glycosyltransferase. Viruses 2022; 14: 1088.

53. Yoon H-S, Golden JW. Heterocyst pattern formation controlled by a diffusible peptide. Science 1998; 282: 935–938.

54. Khudyakov IY, Golden JW. Different functions of HetR, a master regulator of heterocyst differentiation in *Anabaena* sp. PCC 7120, can be separated by mutation. Proc Natl Acad Sci USA 2004; 101: 16040–16045.

55. Wong FCY, Meeks JC. The *hetF* gene product is essential to heterocyst differentiation and affects *hetR* function in the cyanobacterium *Nostoc punctiforme*. J Bacteriol 2001; 183: 2654– 2661.

56. Risser DD, Callahan SM. HetF and PatA control levels of HetR in *Anabaena* sp. strain PCC 7120. J Bacteriol 2008; 190: 7645–7654.

57. Buikema WJ, Haselkorn R. Characterization of a gene controlling heterocyst differentiation in the cyanobacterium *Anabaena* 7120. Genes Dev 1991; 5: 321–330.

58. Videau P, Rivers OS, Hurd K, Ushijima B, Oshiro RT, Ende RJ, et al. The heterocyst regulatory protein HetP and its homologs modulate heterocyst commitment in *Anabaena* sp. strain PCC 7120. Proc Natl Acad Sci U S A 2016; 113: E6984–E6992.

59. Videau P, Rivers OS, Tom SK, Oshiro RT, Ushijima B, Swenson VA, et al. The hetZ gene indirectly regulates heterocyst development at the level of pattern formation in Anabaena sp. strain PCC 7120. 2018; 109: 91–104.

60. Borthakur PB, Orozco CC, Young-Robbins SS, Haselkorn R, Callahan SM. Inactivation of *patS* and *hetN* causes lethal levels of heterocyst differentiation in the filamentous cyanobacterium *Anabaena* sp. PCC 7120. Mol Microbiol 2005; 57: 111–123.

61. Wolk CP, Fan Q, Zhou R, Huang G, Lechno-Yossef S, Kuritz T, et al. Paired cloning vectors for complementation of mutations in the cyanobacterium *Anabaena* sp. strain PCC 7120. Arch Microbiol 2007; 188: 551–563.

62. Xing W-Y, Liu J, Wang Z-Q, Zhang J-Y, Zeng X, Yang Y, et al. HetF protein is a new divisome component in a filamentous and developmental cyanobacterium. mBio 2021; 12: e01382–21.

63. Rodriguez-Valera F, Martin-Cuadrado AB, Rodriguez-Brito B, Pašić L, Thingstad TF, Rohwer F, et al. Explaining microbial population genomics through phage predation. Nat Rev Microbiol 2009; 7: 828–836.

64. Sukenik A, Hadas O, Kaplan A, Quesada A. Invasion of Nostocales (cyanobacteria) to subtropical and temperate freshwater lakes - physiological, regional, and global driving forces. Front Microbiol 2012; 3: 1–9.

65. Stüken A, Rücker J, Endrulat T, Preussel K, Hemm M, Nixdorf B, et al. Distribution of three alien cyanobacterial species (Nostocales) in northeast Germany: *Cylindrospermopsis raciborskii*, *Anabaena bergii* and *Aphanizomenon aphanizomenoides*. Phycologia 2006; 45: 696–703.

66. Pollard PC, Young LM. Lake viruses lyse cyanobacteria, *Cylindrospermopsis raciborskii*, enhances filamentous-host dispersal in Australia. Acta Oecologica 2010; 36: 114–119.

67. Huisman J, Codd GA, Paerl HW, Ibelings BW, Verspagen JMH, Visser PM. Cyanobacterial blooms. Nat Rev Microbiol 2018; 16: 471–483.

68. Alster A, Kaplan-Levy RN, Sukenik A, Zohary T. Morphology and phylogeny of a non-toxic invasive *Cylindrospermopsis raciborskii* from a Mediterranean Lake. Hydrobiologia 2010; 639: 115–128.

69. Brenes-Álvarez M, Vioque A, Muro-Pastor AM. The integrity of the cell wall and its remodeling during heterocyst differentiation are regulated by phylogenetically conserved small RNA Yfr1 in *Nostoc* sp. strain PCC 7120. mBio 2020; 11: e02599–19.

70. Stanier RY, Deruelles J, Rippka R, Herdman M, Waterbury JB. Generic Assignments, Strain Histories and Properties of Pure Cultures of Cyanobacteria. Microbiology 1979; 111: 1–61.

71. Khudiakov II, Gromov B V. Temperate cyanophage A-4 (L) of the blue-green alga *Anabaena variabilis*. Mikrobiologiia 1973; 42: 904–7.

72. Laloum E, Cattan-Tsaushu E, Schwartz DA, Shaalan H, Enav H, Kolan D, et al. Isolation and characterization of a novel Lambda-like phage infecting the bloom-forming cyanobacteria *Cylindrospermopsis raciborskii*. Environ Microbiol 2022; 24: 2435–2448.

73. Holm-Hansen O, Lorenzen CJ, Holmes RW, Strickland JDH. Fluorometric Determination of Chlorophyll. ICES J Mar Sci 1965; 30: 3–15.

74. Zborowsky S, Lindell D. Resistance in marine cyanobacteria differs against specialist and generalist cyanophages. Proc Natl Acad Sci USA 2019; 116: 16899–16908.

75. Munawaroh HSH, Apdila ET, Awai K. *hetN* and *patS* mutations enhance accumulation of fatty alcohols in the *hglT* mutants of *Anabaena* sp. PCC 7120. Front Plant Sci 2020; 11: 1–8.

76. Baym M, Kryazhimskiy S, Lieberman TD, Chung H, Desai MM, Kishony RK. Inexpensive multiplexed library preparation for megabase-sized genomes. PLoS One 2015; 10: 1–15.

77. Deatherage DE, Barrick JE. Identification of mutations in laboratory-evolved microbes from next-generation sequencing data using breseq. Methods in Molecular Biology. 2014. pp 165– 188.

78. Robinson JT, Thorvaldsdóttir H, Winckler W, Guttman M, Lander ES, Getz G, et al. Integrative genomics viewer. Nat Biotechnol 2011; 29: 24–26.

79. Finn RD, Mistry J, Tate J, Coggill P, Heger A, Pollington JE, et al. The Pfam protein families database. Nucleic Acids Res 2009; 38: 211–222.

80. Tatusov RL, Galperin MY, Natale DA, Koonin E V. The COG database: A tool for genome-scale analysis of protein functions and evolution. Nucleic Acids Res 2000; 28: 33–36.

81. Mavromatis K, Chu K, Ivanova N, Hooper SD, Markowitz VM, Kyrpides NC. Gene context analysis in the Integrated Microbial Genomes (IMG) Data Management system. PLoS One 2009; 4: e7979.

82. Yu NY, Wagner JR, Laird MR, Melli G, Rey S, Lo R, et al. PSORTb 3.0: Improved protein subcellular localization prediction with refined localization subcategories and predictive capabilities for all prokaryotes. Bioinformatics 2010; 26: 1608–1615.

83. Huynen M, Snel B, Lathe W, Bork P. Predicting protein function by genomic context: quantitative evaluation and qualitative inferences. Genome Res 2000; 10: 1204–1210.

84. Rogozin IB, Makarova KS, Murvai J, Czabarka E, Wolf YI, Tatusov RL, et al. Connected gene neighborhoods in prokaryotic genomes. Nucleic Acids Res 2002; 30: 2212–2223.

85. Couvin D, Bernheim A, Toffano-Nioche C, Touchon M, Michalik J, Néron B, et al. CRISPRCasFinder, an update of CRISRFinder, includes a portable version, enhanced performance and integrates search for Cas proteins. Nucleic Acids Res 2018; 46: W246–W251.

86. Grissa I, Vergnaud G, Pourcel C. CRISPRFinder: a web tool to identify clustered regularly interspaced short palindromic repeats. Nucleic Acids Res 2007; 35: W52–W57.

87. Solovyev V, Salamov A. Automatic annotation of microbial genomes and metagenomic sequences. In metagenomics and its applications in agriculture, biomedicine and environmental studies (Li RE, ed). Nov Sci Publ 2011; 61–78.

88. Stothard P. The Sequence Manipulation Suite: JavaScript programs for analyzing and formatting protein and DNA sequences. Biotechniques 2000; 28: 1102–1104.

89. Gruber AR, Lorenz R, Bernhart SH, Neubock R, Hofacker IL. The Vienna RNA Websuite. Nucleic Acids Res 2008; 36: W70–W74.

90. Lorenz R, Bernhart SH, Höner zu Siederdissen C, Tafer H, Flamm C, Stadler PF, et al. ViennaRNA Package 2.0. Algorithms Mol Biol 2011; 6: 26.

91. Eren AM, Esen ÖC, Quince C, Vineis JH, Morrison HG, Sogin ML, et al. Anvi’o: an advanced analysis and visualization platform for ‘omics data. PeerJ 2015; 3: e1319.

92. Delmont TO, Eren AM. Linking pangenomes and metagenomes: the *Prochlorococcus* metapangenome. PeerJ 2018; 6: e4320.

93. Camacho C, Coulouris G, Avagyan V, Ma N, Papadopoulos J, Bealer K, et al. BLAST+: Architecture and applications. BMC Bioinformatics 2009; 10: 1–9.

94. Plotly Technologies Inc. https://plotly.com/ Collaborative data science. 2015. Plotly Technologies Inc, Montréal.

