## Supplemental text, tables and figures for "Tradeoffs between phage resistance and nitrogen fixation drive the evolution of genes essential for cyanobacterial heterocyst functionality"

### Supplementary text

#### Isolation and characterization of new *C. raciborskii* phages

In September 2017, 11 phages infecting *C. raciborskii* KLL07 were isolated during a *Cylindrospermopsis* bloom in Lake Kinneret, three of which (Cr-LKS4, Cr-LKS5, and Cr-LKS6) are used in this study. The genomes of the phage isolates were sequenced (see supplementary text) and compared to the GenBank dataset. These phages had genomes with high similarity to each other and all belong to the same species (Table S9). These phages all belong to the same genus as Cylindrospermopsis raciborskii virus RM-2018a isolate CrV-01T (hereafter CrV, which is currently unclassified). Due to the high genomic similarity among these new isolates, the architype, Cr-LKS4, was characterized, and the natural variation among the other two phages is shown in Table S9.

The genome of Cr-LKS4 is 101,125 bp long, and has a GC content of 39.5%, which is nearly identical to that of its host (40.2%). It contains 116 ORFs (Fig. S5, Table S8), 106 of which are homologs of genes in CrV (Table S8). For 102 of the latter genes, the homolog in CrV was the best or the only hit. Four genes (gp_002, gp_094, gp_101, and gp_104) had best hits in *Cylindrospermopsis* strains. However, it is unclear whether the phylogenetic origin of the Cr-LKS4 genes is CrV (suggesting vertical gene transfer) or *Cylindrospermopsis* (suggesting horizontal gene transfer). The last 10 genes were probably acquired by lateral gene transfer from *Cylindrospermopsis* (gp_084 and gp_114; Fig S6), other *cyanobacteria* (gp_034; Fig S6), heterotrophic bacteria (gp_092, gp_093, gp_105, and gp_106; Fig S6), or other unknown sources (gp_045, gp_062, and gp_080; had no homologs in the nr protein database; Table S8).

#### Mutant genes in resistance strains

**RA1:** This strain has an insertion of a single nucleotide in the gene all1304 which encodes for bicarbonate transporter BicA, which is localized in the cytoplasmic membrane (Table S1, S3). To our knowledge, this gene was not reported in the context of nitrogen fixation or resistance to phages in *Nostoc* 7120 before. An identical mutation evolved in parallel in an additional resistant strain (RE7, Table S1, S3) that evolved from a different susceptible WT strain. RE7 had additional mutations and was not part of our phenotypic analysis. An additional resistant strain (RE2) had a different mutation in this gene, in addition to two other mutations in different loci

(Table S1). Moreover, two mutations in this gene were identified in the susceptible WTs (Table S2). However, unlike the mutations in the resistant strains that resulted in a frameshift, the mutations in the WT strains caused a single amino acid change. Therefore, the effect of mutations in this gene on the bacterial function is yet to be understood.

**RA2**: This strain completely lost the ability to induce heterocysts and subsequently collapsed during nitrogen starvation (Fig. S2G2-3). Similarly to RD2 which also could not produce heterocysts, RA2 demonstrates an aberrant cell morphology (Fig. S1B). The *nifH* expression of RA2 48 hours after nitrogen stepdown was significantly lower relative to the susceptible paired control. This strain had two mutations. The first mutation is a SNP in alr3683 that encodes for an ATP-dependent Clp protease proteolytic subunit. In *Synechococcus* sp. PCC 7942, Clp proteases are known to have a role in acclimatization for strong light, cold temperature or UV-B light [1]. The second mutation is a deletion in all5094, which encodes for an oxidoreductase. It is not clear yet how either of these genes affect resistance to phages or heterocysts loss.

**RB1, RB8, RB9, and RB20:** These strains carry a single mutation, causing a frameshift in alr4494 that encodes for glycosyltransferase (Table S1, S3). We performed physiological tests on RB1, RB8, RB20 that were initially selected with A-4L (RB1) or with AN15 (RB8, RB20). These strains showed a significant reduction in growth under nitrogen starvation (Fig. S2 E2-3) and collapsed eventually. The *nifH* expression in these strains was significantly lower relative to the susceptible WT (23% for RB1, 29% for RB8, and 14% for RB20; Fig. S3B). RB1 has narrower cells relative to the susceptible WT (2.56 µm and 3 µm respectively; Fig. S1), with no significant change in the cell length in comparison to the susceptible WT. The heterocysts of this strain appeared in groups of two to three adjacent cells (Fig. S3C) similarly to RE1 (See main text). The gene alr4494 is found two genes upstream of all4496 that encodes for HepK, a protein involved in formation of heterocyst specific glycolipid (HGL) [2, 3]. alr4494 is found in an operon consisting of cell-surface related genes, that was occasionally mutated in the resistant strains (Fig. 5C, Table S1). Three additional strains carry the same mutation (Table S1, strains RB10, RB11, and RB13) and potentially other mutations, and thus were not phenotypically characterized.

**RB2:** This strain had a single mutation in alr4491 that caused a frameshift in a gene encoding an enzyme that has a pyrophosphorylase activity on dTDP-glucose [4] that is involved in cell wall biosynthesis. This gene is located in the same operon as alr4494, however, to the best of our knowledge, it was not reported to be related to nitrogen fixation or heterocyst formation or functionality. The cells of this strain were 15% narrower than those of the WT, yet no significant change in their length was observed (Fig. 1A and S1A). In addition, a synonymous mutation was observed in this gene in the susceptible wild types (Table S2).

**RB4:** Similarly to RB2, this strain had a mutation in alr4491. However, while the mutation in RB2 caused a frameshift, the mutation in RB4 caused an amino acid substitution. As in RB2, cells of RB4 were 15% narrower than those of the WT, yet no significant change in length was observed (Fig. 1A and S1A).

**RD1:** This strain had a single mutation in a gene encoding for a DNA-binding transcriptional regulator (all1719; Table 1, S1). This resistant strain had a reduced heterocyst induction (27%; Table 1) and nearly no nitrogenase gene expression (4%) and activity (3%). This suggests that this gene takes part in the regulation of heterocyst differentiation as well as in nitrogen fixation in the induced heterocyst cells (Fig. 3). Nevertheless, we did not observe an impaired structural appearance of the heterocyst cells (Fig. 3D). Homologs of this gene are found in other members of the *Nostocales,* as well as in other nitrogen fixing *cyanobacteria* and heterotrophic bacteria, however, in a very low similarity, and it is missing from others (Fig. S4). Moreover, this gene clustered, together with a few cyanobacterial genes, with heterotrophic bacteria, while homologs in other *cyanobacteria* strains clustered separately (Fig. S4). This suggests that this gene is an accessory gene acquired by lateral gene transfer. To our knowledge, the relation of this gene to nitrogen fixation and heterocyst induction has not been studied yet.

**RD2:** This strain could not produce heterocysts and had no nitrogenase activity. Moreover, the cells of this strain had an aberrant morphology (Fig. 1B and 3D) both in nitrogen rich and poor environments. It has two SNPs in two loci. The first mutation is intergenic: 518 bp upstream of all1059 and 197 bp upstream of all1060. all1059 encodes for a sucrose synthase, which is involved in disaccharides metabolism. This gene was up-regulated during desiccation and might have a role in the synthesis of osmoprotacnts [5]. A previous study showed that a homolog of this gene in *Anabaena variabilis* is involved in the synthesis of polysaccharides [6], a main component of the bacterial lipopolysaccharide (LPS) that can serve as a receptor for an infection by the phage. Indeed, LPS serves as a receptor for A-4L [3]. all1059 had homologs in all

*Nostocales* strains analyzed in this study, except the *Cylindrospermopsis* and *Raphidiopsis* strains. all1060 encodes for a 150 bp long hypothetical protein, that looks like a *cyanobacteria* specific core gene. The distance of the mutation from both genes (especially from all1059) makes it unclear whether this mutation takes any part in the regulation of these genes, and thus whether it affects any of the phenotypic changes (heterocyst differentiation or phage resistance) of this strain.

The second mutation was found in the gene all2170, which encodes for a Caspase HetF Associated with the Tetratricopeptide repeats (CHAT) domain containing protein. HetF (encoded by alr3546) is an essential regulator of heterocyst differentiation in *Nostoc punctiforme* and in *Nostoc* 7120 [7–9]. Furthermore, HetF also has a role in cell division of *Nostoc* 7120 cells, though it is not essential for division [10]. Both HetF and all2170 belong to the same protein family (pfam12770). The HetF family belongs to the caspase–hemoglobinase fold (CHF) proteases. These CHF proteases can serve as negative regulators in cases where several copies of CHF proteins are expressed, by creating a dimer [11]. Homologs of this gene were found in most of the *cyanobacteria* strains analyzed, similarly to the *hetF* gene. However, while some strains (*Synechococcus* sp. WH 8101, *Synechococcus* sp. JA-2-3B'a(2-13), and *Prochlorococcus marinus* MED4) miss both genes, others have either all2170 (*Cyanothece* sp. UBA12306, *Cyanothece* sp. BG0011, *Lyngbya confervoides* BDU141951, *Pseudanabaena* sp. PCC 7367, and *Pseudanabaena* sp. ABRG5-3) or *hetF* (*C. raciborskii* KLL07, *C. raciborskii* Cr2010, and *Raphidiopsis brookii* D9).

The phenotype of RD2 is similar to that of previously described mutants of *hetF* which might imply a similar function in context of heterocyst differentiation (See main text).

Resistance in this strain was due to a significant reduction (~90%) in the adsorption of the phage to this strain, when compared to the susceptible ancestor. It was previously shown that the O antigen of lipopolysaccharide (LPS) of *Nostoc* 7120 serves as the receptor of A-4L [3]. Multiple genes take part in the construction and modification of the O antigen, and thus a mutation in various loci within these genes can potentially affect the structure of the O antigen and the ability of the phage to identify this receptor. It is thus still unclear which mutation is the resistance conferring mutation in this strain.

**RD3:** This strain had a significantly impaired adsorption of phage particles to the cell surface (Fig. 1C). Additionally, the induction of heterocyst cells as well as the expression of the *nifH* gene, under nitrogen starvation, were lower by 68% and 80% respectively, compared to those of the paired control (Fig. S3). Yet, the heterocysts of this strain had a normal appearance and no tradeoff has been observed during nitrogen starvation (Fig. S2, S3).

This strain carries a single mutation in all1058. This gene encodes for a protein that has two domains: trehalose and maltose phosphorylase, and β -phosphoglucomutase. This gene was found to be up-regulated in *Nostoc* 7120 during desiccation, suggesting that it might has a role in the synthesis of osmoprotectans [5], which are often polysaccharide molecules [12]. β-phosphoglucomutase is a type of isomerase which catalyzes the reaction D-glucose 1-phosphate and D-glucose 6-phosphate. It has been shown that β-phosphoglucomutase formation is induced by the presence of maltose and it is involved in cell wall polysaccharides biosynthesis [13–16]. Modifications in the cell wall polysaccharides may be the cause for the impaired adsorption of the phage A-4L to this strain [3]. It is unclear how this mutation caused a decrease in the ability of strain RD3 to induce heterocyst cells as well as a decrease in *nifH* expression and at the same time did not significantly affect the growth in nitrogen poor medium.

**RD4:** This strain had a significant reduction in growth in comparison to its susceptible paired control, however, while it was able to grow in nitrogen rich medium, its growth was almost arrested under nitrogen starvation (Fig. S2). The phenotypic appearance of the heterocysts showed the lack of cyanophycin polar granules (Fig. S3) which is a nitrogen reserve of fixed nitrogen, that can be transferred to the vegetative cells in the form of arginine and aspartate [17, 18]. This might suggest that impaired nitrogen fixation activity of RD4 leads to the reduced growth under nitrogen deprivation. We identified in RD4 a single mutation in the gene alr4485, which encodes for an ABC-type polysaccharide/polyol phosphate export permease. This transporter is embedded in the inner membrane in gram negative bacteria, and exports a variety of substrates, among them, cell wall components, such as polysaccharides, and the O-antigenic polysaccharide [19, 20]. Therefore, a mutation in this gene may affect the cell wall composition and thus cause impaired phage adsorption. A previous study showed that the gene *hetC* (alr2817), which has a role in the regulation of differentiation to heterocysts, is similar to bacterial ABC membrane transport proteins [21], though aligning the sequences of *hetC* and alr4485 showed no significant similarity between them (data not shown).

**RD5:** This strain had reduced growth under nitrogen starvation conditions relative to the susceptible paired control (Fig. S2). Bright field image of RD5 suggests structurally normal heterocyst cells (Fig. S3). We identified SNP in a gene (alr1906) predicted to be a member of the FhaB family (Filamentous hemagglutinin), which is a part of a two-partner secretion system (Tps). This protein (TpsA) is transported by a transporter protein (TpsB) from the periplasmic side and secreted to the extracellular medium, in various gram-negative bacteria [22]. A study that examined two TpsB-like proteins in *Nostoc* 7120, suggested that alr1906 is one of the substrates of the studied TpsB-like proteins [23]. Under nitrogen starvation, strains carrying mutations in the TpsB-like proteins, showed structurally normal heterocysts, yet a reduced growth relative to the control strain was noticed. Moreover, this disruption was accompanied by resistance to antibiotics (erythromycin, roxithromycin, and tylosin).

**RE1:** This strain had a reduced ability to induce heterocyst cells, however, this reduction was limited in comparison to other resistant strains (Fig. 3B). While this strain had only 27% less heterocysts in comparison to the wild type, it had a very low *nifH* expression and no nitrogenase activity (Fig. 3), and it could not survive nitrogen starvation (Fig. 2B). The heterocysts of this strain had a deformed morphology and sometimes appeared in groups of 2-3 adjacent cells in different stages of differentiation (Fig. 3D), which may explain their malfunction. Furthermore, this strain had a significantly impaired adsorption of phage A-4L. RE1 has a mutation in the all5019, which encodes for 3-phosphoshikimate 1-carboxyvinyltransferase that is involved in aromatic amino acid biosynthesis (Powell et al. 1992). Phylogeny of this gene suggests that this gene was acquired by lateral gene transfer (Fig. S4) as has been shown previously using phylogeny, structure and activity of this enzyme [24]. We identified the same mutation in the substrain RE8 (Table S1), which originated from the same susceptible ancestor (WT-E). However, the mutation in RE8 was one of five mutations, which suggests that RE8 evolved from RE1. To the best of our knowledge, this gene was not linked previously to nitrogen fixation of heterocyst differentiation or to resistance to phages.

**RE3:** This strain could not survive nitrogen starvation (Fig. S2) and was able to induce heterocyst cells with a similar phenotype to RE1. We found in this strain a mutation in an intergenic region upstream to all3346 that encodes for a Repeats-in-toxin (RTX) toxin-related $Ca^{2+}$-binding protein, and downstream to all3347 that encodes for a CHASE2 domain-containing protein. RTX proteins are a family of proteins, which are secreted via a type I secretion system. RTX proteins have various functions in gram-negative bacteria. In unicellular *cyanobacteria* RTX proteins are related to cellular motility (Linhartová et al. 2010, Baumann 2019). Hahn et al. (2015) identified three possible members of the RTX proteins family in *Nostoc* 7120 (all0275, alr4238 and all7128) but their function remains unclear. Though the mutation position of RE3 was not predicted to be a promoter site, the morphological structure of its heterocysts suggests a regulatory role of this locus which is common for non-coding regions.

**RE4:** This strain produced significantly less (60%) heterocysts than the WT (Fig S3). Although this strain produces heterocysts that appear to be structurally normal, the population of this strain had reduced growth under nitrogen starvation (Fig. S2), followed by drastically low *nifH* expression (Fig. S3). Moreover, our experiments indicate that the resistance of this strain to A-4L was due to significantly impaired adsorption of phage particles (Fig 1C). This strain had a SNP in the gene all7617, which is located on the beta plasmid of *Nostoc* 7120 (Tables S1, S3) and encodes for a CusA/CzcA family heavy metal efflux RND transporter. A previous study showed that all7617 was upregulated under nitrogen starvation similarly to its two homologs in the *Nostoc* 7120 genome: all7618 and all763 (Flaherty *et al.*, 2011, suppotrting information). These two genes are also located in the beta plasmid and encode for CusA/CzcA family heavy metal efflux RND transporters.

**RE5:** This strain can induce heterocysts, however, it collapsed during nitrogen starvation (Fig. S2). This strain has a synonymous mutation in alr5237 that encodes for glycosyltransferase that is involved in the synthesis of molecules such as polysaccharides that can affect the structure of the cell surface (Lairson et al. 2008). This SNP is positioned on the $30^{th}$ amino acid (leucin, TTA became TTG). Though silent mutations result in the same amino acid, such mutations can affect protein folding [26], which can be crucial for enzyme activity. When analyzing the main chromosome to understand the codon usage in *Nostoc* 7120, TTA and TTG have almost equal frequency throughout the genome (0.21 and 0.22 respectively). However, alr5237 itself has a different codon usage for leucine, which is compatible with the fact that this gene was acquired by lateral gene transfer (Fig. S4). TTA and TTG have usage frequency of 0.47 and 0.14

respectively, which indicates a codon usage bias. We did not observe any difference between the mRNA secondary structure in silico when replacing the $30^{th}$ codon (TTA) with TTG. When scanning the genome for tRNAs, the frequency for TTA is 0.125 while for TTG it is 0.25. Such differences may affect the translation of this gene. Another resistant substrain (RE6) that belongs to the same lineage, had the same mutation (Table S1). Since RE6 had two additional mutations, which suggests that RE6 evolved from RE5.

**RE2**: The heterocyst induction of RE2 was lower by 65% (Fig. 3B) than that of the wild type strain. Although this strain could produce structurally normal heterocyst cells (Fig. 3D), no nitrogenase activity was observed (Fig. 3A), the expression of the *nifH* gene was significantly lower (Fig. 3D), and it could not survive nitrogen starvation (Fig. 2B).

Strain RE2 has three mutations (Table S1). The first mutation is a single base deletion in the gene alr4949, which has two conserved domains: a serine/threonine protein kinase (residues ~20-270) and a porphyrin-binding protein domain GUN4 (residues ~290-410). The mutation was detected in the second domain. GUN4 is involved in chlorophyll biosynthesis regulation and intracellular signaling in photosynthetic organisms such as plants and algae [27, 28]. Though, previous studies on green algae reported reduced chlorophyll biosynthesis followed by pale green color of the GUN4 mutants [29, 30], we did not observe remarkable color change of RE2 compared to the susceptible paired control.

An additional mutation was identified in all1304 (deletion of one nucleotide), which encodes for a bicarbonate transporter (BicA). We identified different mutations in this gene both in the susceptible controls and in an additional resistant strain that was mentioned previously (See RA8).

The third mutation in this strain was in the gene alr4492, which encodes for a glycosyltransferase family 2 (GT2) protein. As mentioned above, glycosyltransferases are involved in the synthesis of polysaccharides, which may affect the cell surface structure and thus the ability of phages particles to adsorb to the cell. We identified a mutation in this gene in the strain RF16 (among another three mutations) which was not part of our phenotypic analysis. This gene is part of an operon enriched with cell surface related genes that were found to be essential for phage infection in other strains isolated in this study.

#### Supplementary materials and methods

**Whole genome sequencing of the phages:** Following Laloum et al. (2022), DNA was extracted using the phenol/chloroform method using the protocol in Pickard (2009). Libraries were prepared using Quantabio's sparQ DNA Frag & Library Prep Kit with Illumina's TrueSeq DNA CD indexes. The genome was sequenced on a MiSeq System (Illumina; 2 x 150 bp). A quality assessment of the sequences was performed using FastQC (standard parameters) [32]. Quality trimming was performed using Trim Galore 0.4.2 (parameters: phred 33, quality 20), in order to discard the adapter sequences, and the read quality was reassessed. Assembly was performed using SPAdes 3.9.0 (phred offset 33, threads 24, only-assembler, K-127, 77, 55,105) [33]. The assembly resulted in three scaffolds per genome. In each genome, two scaffolds had a similar coverage, while the third scaffold had approximately double coverage (Table S6). A similar phenomenon was described for the genome of Cylindrospermopsis raciborskii virus RM-2018a isolate CrV-01T [34]. It was discovered that the genome of this phage contains a large inverse duplication that prevented the full assembly of the genome. The DNA sequence of our phages showed high similarity to this phage, suggesting that our genomes contain inverse duplication as well. To verify that this is the case in our genomes, PCR reactions were performed, using primers designed according to the sequence of the scaffold edges (Table S7). The results of the PCR reactions suggested that, similarly to CrV, the genome sequence of the Cr-LKS4 is circular and contained a reverse duplication. The scaffold order is: Scaffold 1, Scaffold 2 (forward), Scaffold 3, and scaffold 2 (reverse), which is connected to the beginning of scaffold 1. Sanger sequencing of the PCR amplicons enabled the curation of the genome sequence around the connections.

Gene prediction was performed using GeneMarkS with standard parameters [35]. The predicted genes were annotated using blastp [36] against the NCBI non-redundant protein database. Predicted sequences with e-values <$10^{-4}$ were considered to be homologs [34]. In addition, predicted function was assessed using HHpred with default parameters [37]. The database used for these predictions was UniProt-SwissProt-viral70, 3 November 2021. HHpred Probability was indicated (Table S8). Results with probability lower than 95% are not shown. Final gene prediction was assigned considering homologs function, conserved domains suggested by blast, and HHpred prediction. Two CRISPR arrays with evidence level greater than 1 were identified by CRISPRCasFinder [38]. Five short (43-65 amino acids) predicted ORFs that had no similarity to any protein sequence in the n/r protein database, overlapped the CRISPR arrays, and thus they were removed.

**Phylogeny of Cr-LKS4 genes:** Homologs of Cr-LKS4 genes were recruited using blastp [39] against NCBI non-redundant protein database with e-value < $10^{-4}$. For each gene, the first 30 hits were used to construct the initial tree, which was then manually curated, and non-informative nodes were removed.

Multiple sequence alignments were carried using MAFFT online services with default parameters [40], and sequences that did not align well were manually removed. Phylogenetic trees were generated using CIPRES Science Gateway V 3.3 [41] with RAxML Black Box using protein substitution matrix: LG. Bootstraps values were achieved by resampling 100 maximum likelihood trees. Trees were visualized by FigTree v.1.4.4 (http://tree.bio.ed.ac.uk/).

#### Supplementary tables

**Table S4:** Resistant substrains of *C. raciborskii*.

| **Resistant substrain** | **Selecting phage** | [1]**Selection method** |
|---|---|---|
| R3C1 | Cr-LKS4 | Semi-solid medium |
| R3C2 | Cr-LKS5 | Semi-solid medium |
| R3C3 | Cr-LKS6 | Semi-solid medium |
| R3C4 | Cr-LKS4 | Liquid medium |
| R3C5 | Cr-LKS5 | Liquid medium |
| R8D1 | Cr-LKS4 | Liquid medium |
| R8D2 | Cr-LKS5 | Liquid medium |
| R8D3 | Cr-LKS6 | Liquid medium |

[1]S refers to selection in semi-solid medium and L refers to selection in liquid medium.

**Table S5:** Organisms used in the heatmap, pangenome analysis, and phylogenetic analysis.

| [1]**Assembly accession** | [2]**Organism** |
|---|---|
| GCF_000317695.1 | *Anabaena cylindrica* PCC 7122 |
| GCF_000312705.1 | *Anabaena* sp. 90 |
| GCF_001277295.1 | *Anabaena* sp. WA102 |
| GCA_014698355.1 | *Anabaena variabilis* FACHB-164 |
| GCF_002368175.1 | *Calothrix* sp. NIES-2098 |
| GCF_019977735.1 | *Calothrix* sp. PCC 7716 |
| GCF_003013815.1 | *Cyanothece* sp. BG0011 |
| GCA_003448685.1 | *Cyanothece* sp. UBA12306 |
| GCF_003367075.2 | *Cylindrospermopsis raciborskii* Cr2010 |
| GCF_021650815.1 | *Cylindrospermopsis raciborskii* KLL07 |
| GCF_000005845.2 | *Escherichia coli* str. K-12 substr. MG1655 |
| GCF_000317205.1 | *Fischerella muscicola* PCC 7414 |
| GCF_001548455.1 | *Fischerella* sp. NIES-3754 |
| GCF_022372495.1 | *Leptolyngbya boryana* IU 594 |
| GCF_002368255.1 | *Leptolyngbya boryana* NIES-2135 |
| GCF_000173555.1 | *Limnospira maxima* CS-328 |
| GCF_016026175.1 | *Listeria welshimeri* |
| GCF_000817775.3 | *Lyngbya confervoides* BDU141951 |
| GCF_000169095.1 | *Lyngbya* sp. PCC 8106 |
| GCF_000513915.1 | *Mesorhizobium* sp. WSM2561 |
| GCF_009792235.1 | *Microcystis aeruginosa* FD4 |
| GCF_000010625.1 | *Microcystis aeruginosa* NIES-843 |
| GCF_019704275.1 | *Microcystis aeruginosa* NIES-88 |
| GCF_001767235.1 | *Moorena producens* PAL-8-15-08-1 |
| GCF_001007935.1 | *Nitrosomonas communis* Nm2 |
| GCF_001273775.1 | *Nitrospira moscoviensis* NSP M-1 |
| GCF_022376295.1 | *Nodularia sphaerocarpa* UHCC 0038 |
| GCF_003054475.1 | *Nodularia spumigena* UHCC 0039 |
| GCA_000009705.1 | *Nostoc* sp. pcc 7120 |
| GCF_000020025.1 | *Nostoc punctiforme* PCC 73102 |
| GCF_009873495.1 | *Nostoc* sp. ATCC 53789 |
| GCF_003443655.1 | *Nostoc sphaeroides* Kutzing En |
| GCF_000317105.1 | *Oscillatoria acuminata* PCC 6304 |
| GCF_000317475.1 | *Oscillatoria nigro-viridis* PCC 7112 |
| GCF_023983615.1 | *Phormidium yuhuli* AB48 |
| GCF_000011465.1 | *Prochlorococcus marinus* subsp. pastoris str. CCMP1986 MED4 |
| GCF_003967015.1 | *Pseudanabaena* sp. ABRG5-3 |
| GCF_000317065.1 | *Pseudanabaena* sp. PCC 7367 |
| GCA_000175855.1 | *Raphidiopsis brookii* D9 |

| | |
|---|---|
| GCF_011604545.1 | *Rhizobium leguminosarum* bv. Trifolii 3B |
| GCF_012241395.2 | *Rhizobium phaseoli* S3 |
| GCF_009664245.1 | *Sinorhizobium meliloti* AK21 |
| GCF_000013225.1 | *Synechococcus* sp. JA-2-3B'a(2-13) |
| GCF_004209775.1 | *Synechococcus* sp. WH 8101 |
| GCA_018457135.1 | *Trichodesmium erythraeum* GBRTRLIN201 |
| GCF_009856605.1 | *Trichormus variabilis* 0441 |
| GCF_000204075.1 | *Trichormus variabilis* ATCC 29413 |
| GCF_002291405.1 | *Variovorax boronicumulans* J1 |

**Table S6:** Initial assembly of phage genomes.

| Phage | Scaffold 1 | | Scaffold 2 | | Scaffold 3 | |
|---|---|---|---|---|---|---|
| | Length (bp) | Average coverage | Length (bp) | Average coverage | Length (bp) | Average coverage |
| Cr-LKS4 | 73,098 | 65.97 | 10,709 | 186.22 | 5,731 | 93.29 |
| Cr-LKS5 | 73,850 | 133.28 | 11,003 | 246.38 | 5,731 | 125.15 |
| Cr-LKS6 | 74,086 | 103.88 | 10,719 | 195.83 | 5,731 | 99.37 |

**Table S7:** Primers used to connect the phage scaffolds.

| Prime name | Sequence |
|---|---|
| Scaffold 1F | TGTAAATCCCGACCCAAGTAAGTC |
| Scaffold 1R | GGAGAGATTTACGCCCGTCACA |
| Scaffold 2F | AGGCACTCCGCAATTATCTGGGC |
| Scaffold 2R | CTTACTCAGGTCGGGTTAGATTAGAC |
| Scaffold 3F | GACTTCTATTAGCAGGAATGACACAA |
| Scaffold 3R | AGGGTCTTCAGATGCTCTTTGAGA |

[1]GCA and GCF represent data taken from GenBank and RefSeq respectively.
[2]The full names of the organisms used in the heatmap, pangenome analysis, and phylogenetic analysis.

**Table S9:** Genetic variation between the phages that infect *C. raciborskii*.

| Locus in Cr-LKS4 | | Variation in other phages | |
|---|---|---|---|
| Locus | Gene | Cr-LKS5 | Cr-LKS6 |
| 1-7 | Upstream to gp116 (hypothetical protein) | • | TGTCCGT→AGGTC |
| 12,072 | gp013 (hypothetical protein) | G→A | • |
| 40,411 | gp045 (hypothetical protein) | +CATGGGAACATAACTT | +CATGGGAACATAACTT |
| 74,263 | gp082 (hypothetical protein) | C→T | • |
| 80,924-81,673 | gp092 (DnaJ domain-containing protein) | • | Δ750 bp |
| 93,427-94,176 | gp106 (DnaJ domain-containing protein) | • | Δ750 bp |
| 100,837 | gp116 | G→A | • |

#### Supplementary figures

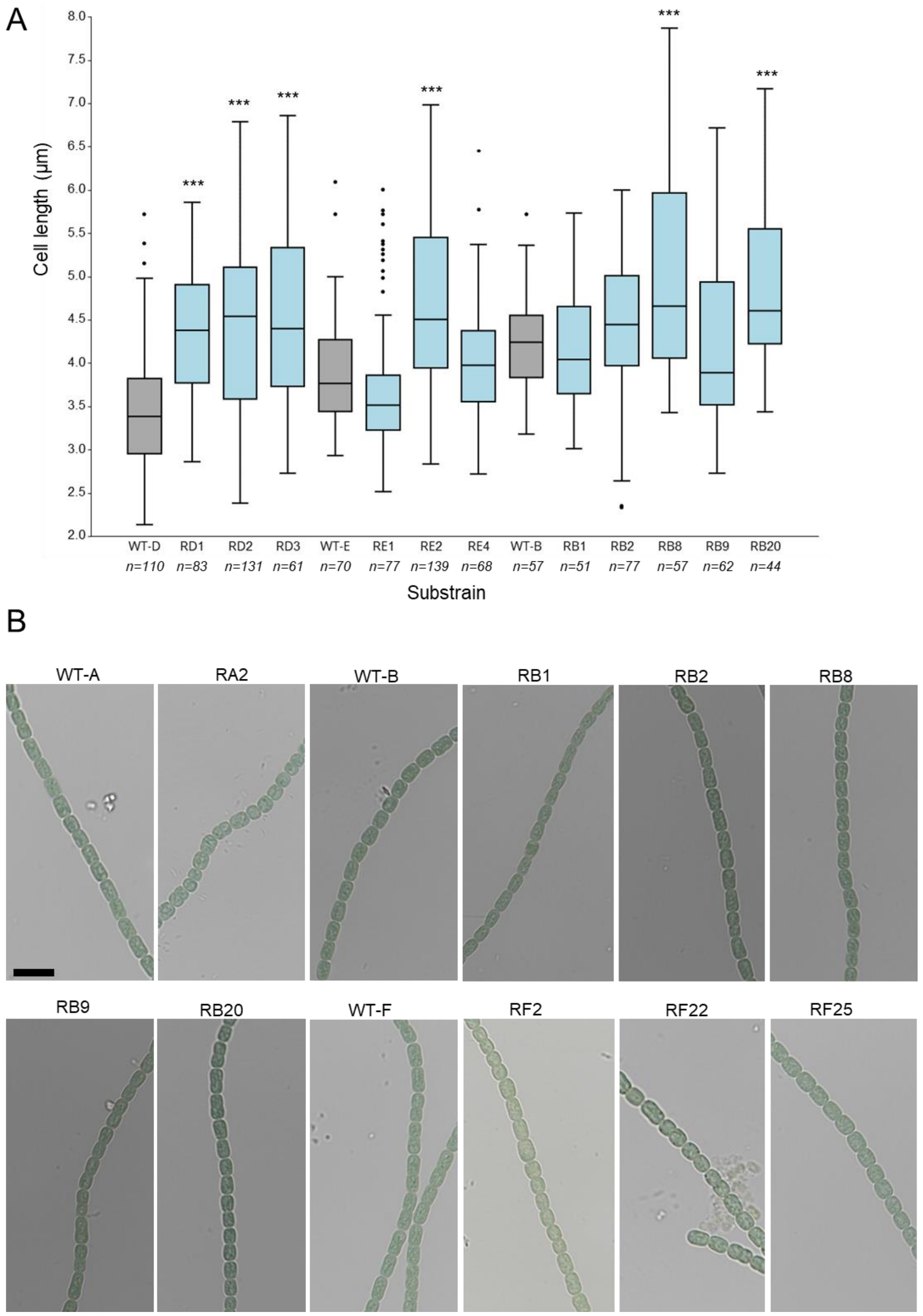


**Figure S1: Phenotypes of the resistant substrains of *Nostoc* 7120.** Cell (**A**) length of the resistant substrains (light blue) and their susceptible ancestors (grey). Data shown are average +/- standard deviation of n replicates. *** $p<0.001$.**B**. Bright Field images of resistant substrains and their susceptible ancestors. Scale equals 10µm.

A1 A2

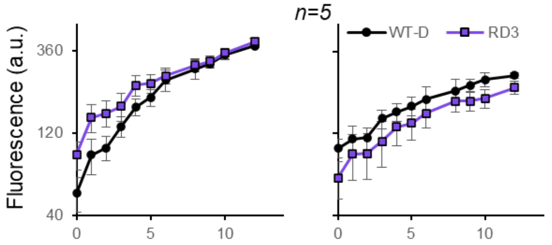


A3

| | +N | -N | -N vs. +N |
|---|---|---|---|
| RD3 | ** | NS | NS |

B1 B2

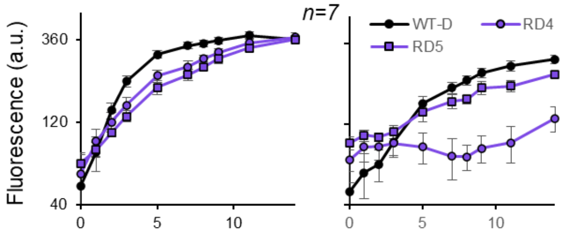


B3

| | +N | -N | -N vs. +N |
|---|---|---|---|
| RD4 | *** | *** | *** |
| RD5 | *** | *** | *** |

C1 C2

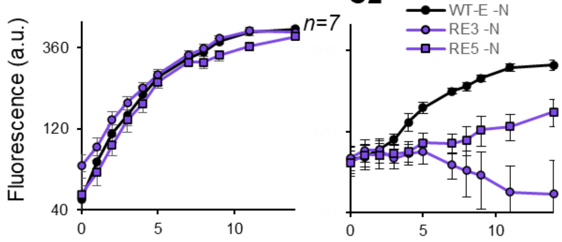


C3

| | +N | -N | -N vs. +N |
|---|---|---|---|
| RE3 | NS | *** | *** |
| RE5 | *** | *** | ** |

D1 D2

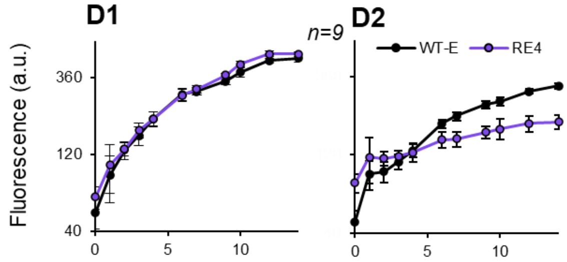


D3

| | +N | -N | -N vs. +N |
|---|---|---|---|
| RE4 | NS | *** | *** |

E1 E2

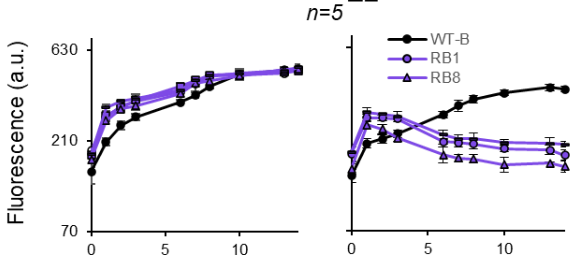


E3

| | +N | -N | -N vs. +N |
|---|---|---|---|
| RB1 | ** | *** | *** |
| RB8 | NS | *** | *** |
| RB20 | NS | *** | *** |

F1 F2

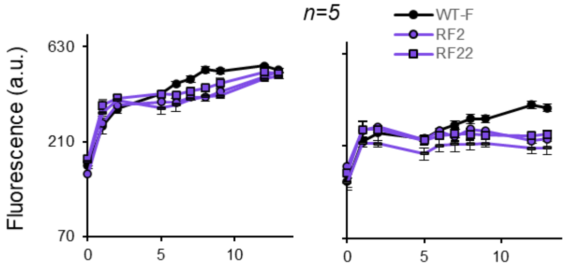


F3

| | +N | -N | -N vs. +N |
|---|---|---|---|
| RF2 | *** | *** | *** |
| RF22 | *** | *** | *** |
| RF25 | NS | *** | *** |

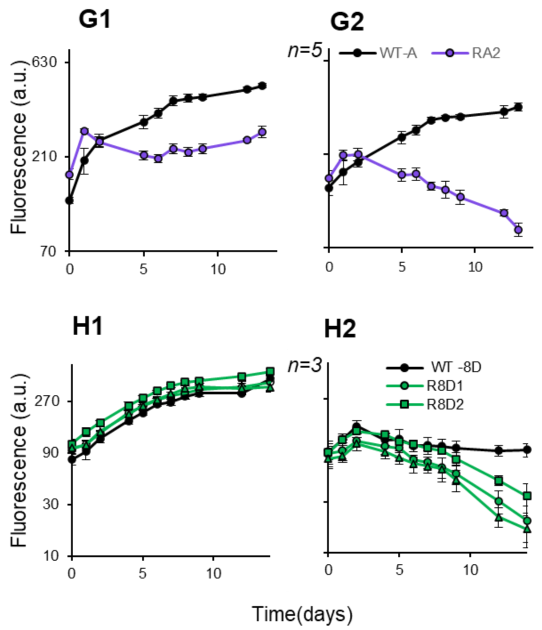


G3

| | +N | -N | -N vs. +N |
|---|---|---|---|
| RA2 | ** | *** | *** |

H3

| | +N | -N | -N vs. +N |
|---|---|---|---|
| R8D1 | * | *** | * |
| R8D2 | NS | *** | *** |
| R8D3 | ** | *** | *** |

**Figure S2: Growth cost of resistant substrains.** Growth of resistnat substrains (purple for *Nostoc* 7120, green for *C. raciborskii*) and wild types (WT; black) and their suceptible paired controls. Growth was measured as chlorophyll *a* autofluorescece with (left panel) or without (middle panel) combined nitrogen. The data presented are average and standard deviation (+/-) of n biological replicates. The corresponding statistical significance of the difference in the growth of the resistant substrains and their susceptible paired controls with (+N) or without (-N) nitrogen as well as the cost in the presence vs. absence of nitrogen (+N vs. -N) are presented in the tables on the right panels (NS non-significant, * $p<0.05$, ** $p<0.01$, *** $p<0.001$). *p*-values were calculated using two-way repeated measure ANOVA tests.

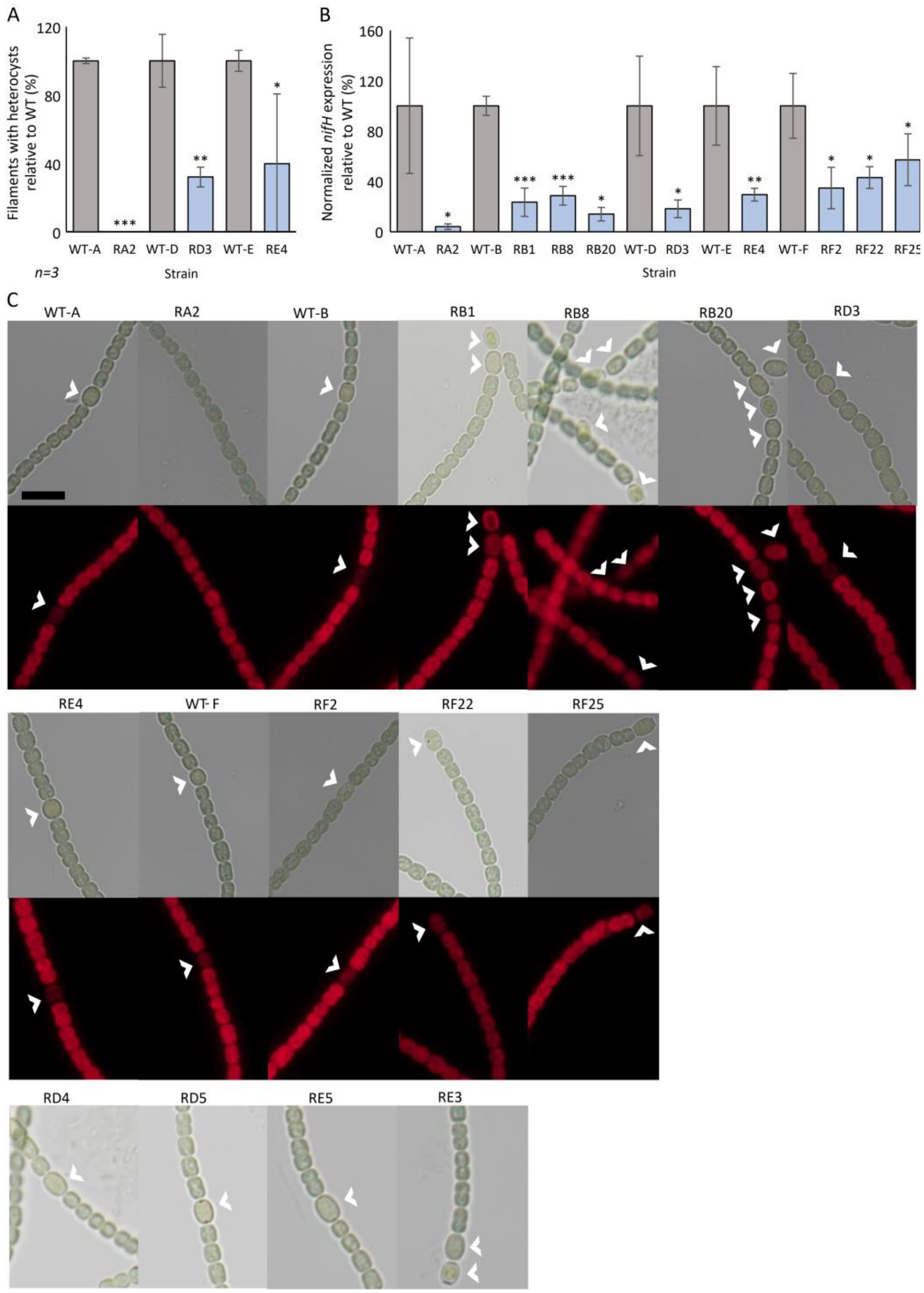


**Figure S3**: **Cost of resistance in nitrogen fixation. A.** Percentage of filaments with heterocysts of the resistant substrains (light blue) relative to their susceptible ancestor (WT, grey), 48 hours after nitrogen stepdown. **B.** Expression of *nifH* gene in the resistant substrains (light blue) relative to their susceptible ancestor (WT, grey), 48 hours after nitrogen stepdown. The transcript levels of *nifH* values are normalized to transcript levels of *rnpB*. Data shown are average and standard deviation (+/-) of n biological replicates. *$p<0.05$, **$p<0.01$, ***$p<0.001$. **C.** Bright field images and the corresponding fluorescence images of *Nostoc* 7120 substrains, 48 hours after nitrogen stepdown. White arrows indicate heterocyst cells. Scale equals 10 µm.

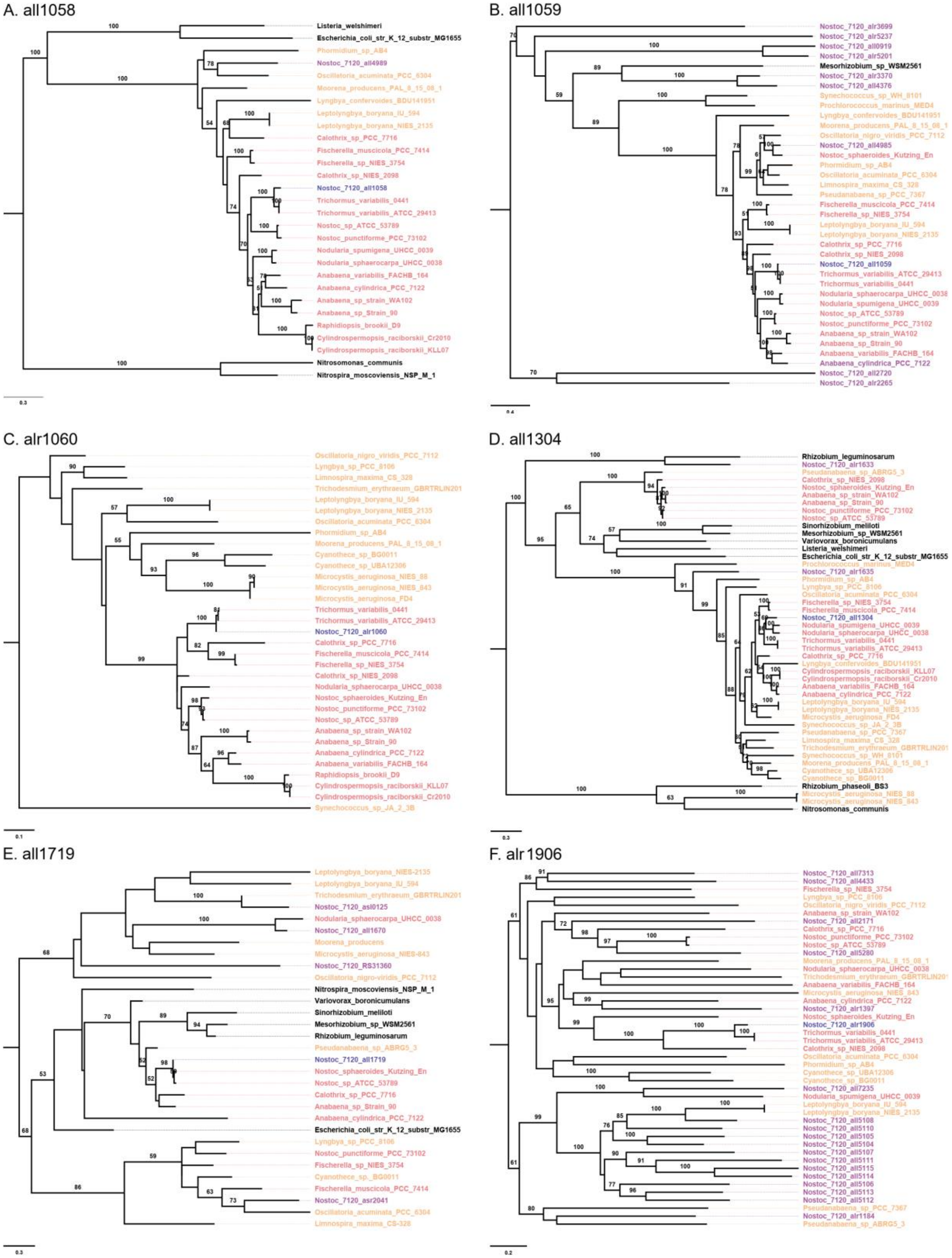
A. all1058
Listeria_welshimeri
Escherichia_coli_str_K_12_substr_MG1655
Phormidium_sp_AB4
Nostoc_7120_all4989
Oscillatoria_acuminata_PCC_6304
Moorena_producens_PAL_8_15_08_1
Lyngbya_confervoides_BDU141951
Leptolyngbya_boryana_IU_594
Leptolyngbya_boryana_NIES_2135
Calothrix_sp_PCC_7716
Fischerella_muscicola_PCC_7414
Fischerella_sp_NIES_3754
Calothrix_sp_NIES_2098
Nostoc_7120_all1058
Trichormus_variabilis_0441
Trichormus_variabilis_ATCC_29413
Nostoc_sp_ATCC_53789
Nostoc_punctiforme_PCC_73102
Nodularia_spumigena_UHCC_0039
Nodularia_sphaerocarpa_UHCC_0038
Anabaena_variabilis_FACHB_164
Anabaena_cylindrica_PCC_7122
Anabaena_sp_strain_WA102
Anabaena_sp_Strain_90
Raphidiopsis_brookii_D9
Cylindrospermopsis_raciborskii_Cr2010
Cylindrospermopsis_raciborskii_KLL07
Nitrosomonas_communis
Nitrospira_moscoviensis_NSP_M_1
B. all1059
Nostoc_7120_alr3699
Nostoc_7120_alr5237
Nostoc_7120_all0919
Nostoc_7120_alr5201
Mesorhizobium_sp_WSM2561
Nostoc_7120_alr3370
Nostoc_7120_all4376
Synechococcus_sp_WH_8101
Prochlorococcus_marinus_MED4
Lyngbya_confervoides_BDU141951
Moorena_producens_PAL_8_15_08_1
Oscillatoria_nigro_viridis_PCC_7112
Nostoc_7120_all4985
Nostoc_sphaeroides_Kutzing_En
Phormidium_sp_AB4
Oscillatoria_acuminata_PCC_6304
Limnospira_maxima_CS_328
Pseudanabaena_sp_PCC_7367
Fischerella_muscicola_PCC_7414
Fischerella_sp_NIES_3754
Leptolyngbya_boryana_IU_594
Leptolyngbya_boryana_NIES_2135
Calothrix_sp_PCC_7716
Calothrix_sp_NIES_2098
Nostoc_7120_all1059
Trichormus_variabilis_ATCC_29413
Trichormus_variabilis_0441
Nodularia_sphaerocarpa_UHCC_0038
Nodularia_spumigena_UHCC_0039
Nostoc_sp_ATCC_53789
Nostoc_punctiforme_PCC_73102
Anabaena_sp_strain_WA102
Anabaena_sp_Strain_90
Anabaena_variabilis_FACHB_164
Anabaena_cylindrica_PCC_7122
Nostoc_7120_all2720
Nostoc_7120_alr2265
C. alr1060
Oscillatoria_nigro_viridis_PCC_7112
Lyngbya_sp_PCC_8106
Limnospira_maxima_CS_328
Trichodesmium_erythraeum_GBRTRLIN201
Leptolyngbya_boryana_IU_594
Leptolyngbya_boryana_NIES_2135
Oscillatoria_acuminata_PCC_6304
Phormidium_sp_AB4
Moorena_producens_PAL_8_15_08_1
Cyanothece_sp_BG0011
Cyanothece_sp_UBA12306
Microcystis_aeruginosa_NIES_88
Microcystis_aeruginosa_NIES_843
Microcystis_aeruginosa_FD4
Trichormus_variabilis_0441
Trichormus_variabilis_ATCC_29413
Nostoc_7120_alr1060
Calothrix_sp_PCC_7716
Fischerella_muscicola_PCC_7414
Fischerella_sp_NIES_3754
Calothrix_sp_NIES_2098
Nodularia_sphaerocarpa_UHCC_0038
Nostoc_sphaeroides_Kutzing_En
Nostoc_punctiforme_PCC_73102
Nostoc_sp_ATCC_53789
Anabaena_sp_strain_WA102
Anabaena_sp_Strain_90
Anabaena_cylindrica_PCC_7122
Anabaena_variabilis_FACHB_164
Raphidiopsis_brookii_D9
Cylindrospermopsis_raciborskii_KLL07
Cylindrospermopsis_raciborskii_Cr2010
Synechococcus_sp_JA_2_3B
D. all1304
Rhizobium_leguminosarum
Nostoc_7120_alr1633
Pseudanabaena_sp_ABRG5_3
Calothrix_sp_NIES_2098
Nostoc_sphaeroides_Kutzing_En
Anabaena_sp_strain_WA102
Anabaena_sp_Strain_90
Nostoc_punctiforme_PCC_73102
Nostoc_sp_ATCC_53789
Sinorhizobium_meliloti
Mesorhizobium_sp_WSM2561
Variovorax_boronicumulans
Listeria_welshimeri
Escherichia_coli_str_K_12_substr_MG1655
Prochlorococcus_marinus_MED4
Nostoc_7120_alr1635
Phormidium_sp_AB4
Lyngbya_sp_PCC_8106
Oscillatoria_acuminata_PCC_6304
Fischerella_sp_NIES_3754
Fischerella_muscicola_PCC_7414
Nostoc_7120_all1304
Nodularia_spumigena_UHCC_0039
Nodularia_sphaerocarpa_UHCC_0038
Trichormus_variabilis_0441
Trichormus_variabilis_ATCC_29413
Calothrix_sp_PCC_7716
Lyngbya_confervoides_BDU141951
Cylindrospermopsis_raciborskii_KLL07
Cylindrospermopsis_raciborskii_Cr2010
Anabaena_variabilis_FACHB_164
Anabaena_cylindrica_PCC_7122
Leptolyngbya_boryana_IU_594
Leptolyngbya_boryana_NIES_2135
Microcystis_aeruginosa_FD4
Synechococcus_sp_JA_2_3B
Pseudanabaena_sp_PCC_7367
Limnospira_maxima_CS_328
Trichodesmium_erythraeum_GBRTRLIN201
Synechococcus_sp_WH_8101
Moorena_producens_PAL_8_15_08_1
Cyanothece_sp_UBA12306
Cyanothece_sp_BG0011
Rhizobium_phaseoli_BS3
Microcystis_aeruginosa_NIES_88
Microcystis_aeruginosa_NIES_843
Nitrosomonas_communis
E. all1719
Leptolyngbya_boryana_NIES-2135
Leptolyngbya_boryana_IU_594
Trichodesmium_erythraeum_GBRTRLIN201
Nostoc_7120_asl0125
Nodularia_sphaerocarpa_UHCC_0038
Nostoc_7120_all1670
Moorena_producens
Microcystis_aeruginosa_NIES-843
Nostoc_7120_RS31360
Oscillatoria_nigro-viridis_PCC_7112
Nitrospira_moscoviensis_NSP_M_1
Variovorax_boronicumulans
Sinorhizobium_meliloti
Mesorhizobium_sp_WSM2561
Rhizobium_leguminosarum
Pseudanabaena_sp_ABRG5_3
Nostoc_7120_all1719
Nostoc_sphaeroides_Kutzing_En
Nostoc_sp_ATCC_53789
Calothrix_sp_PCC_7716
Anabaena_sp_Strain_90
Anabaena_cylindrica_PCC_7122
Escherichia_coli_str_K_12_substr_MG1655
Lyngbya_sp_PCC_8106
Nostoc_punctiforme_PCC_73102
Fischerella_sp_NIES_3754
Cyanothece_sp_BG0011
Fischerella_muscicola_PCC_7414
Nostoc_7120_asr2041
Oscillatoria_acuminata_PCC_6304
Limnospira_maxima_CS-328
F. alr1906
Nostoc_7120_all7313
Nostoc_7120_all4433
Fischerella_sp_NIES_3754
Lyngbya_sp_PCC_8106
Oscillatoria_nigro_viridis_PCC_7112
Anabaena_sp_strain_WA102
Nostoc_7120_all2171
Calothrix_sp_PCC_7716
Nostoc_punctiforme_PCC_73102
Nostoc_sp_ATCC_53789
Nostoc_7120_all5280
Moorena_producens_PAL_8_15_08_1
Nodularia_sphaerocarpa_UHCC_0038
Trichodesmium_erythraeum_GBRTRLIN201
Anabaena_variabilis_FACHB_164
Microcystis_aeruginosa_NIES_843
Anabaena_cylindrica_PCC_7122
Nostoc_7120_alr1397
Nostoc_sphaeroides_Kutzing_En
Nostoc_7120_alr1906
Trichormus_variabilis_0441
Trichormus_variabilis_ATCC_29413
Calothrix_sp_NIES_2098
Oscillatoria_acuminata_PCC_6304
Phormidium_sp_AB4
Cyanothece_sp_UBA12306
Cyanothece_sp_BG0011
Nostoc_7120_all7235
Nodularia_spumigena_UHCC_0039
Leptolyngbya_boryana_IU_594
Leptolyngbya_boryana_NIES_2135
Nostoc_7120_all5108
Nostoc_7120_all5110
Nostoc_7120_all5105
Nostoc_7120_all5104
Nostoc_7120_all5107
Nostoc_7120_all5111
Nostoc_7120_all5115
Nostoc_7120_all5114
Nostoc_7120_all5106
Nostoc_7120_all5113
Nostoc_7120_all5112
Pseudanabaena_sp_PCC_7367
Nostoc_7120_alr1184
Pseudanabaena_sp_ABRG5_3

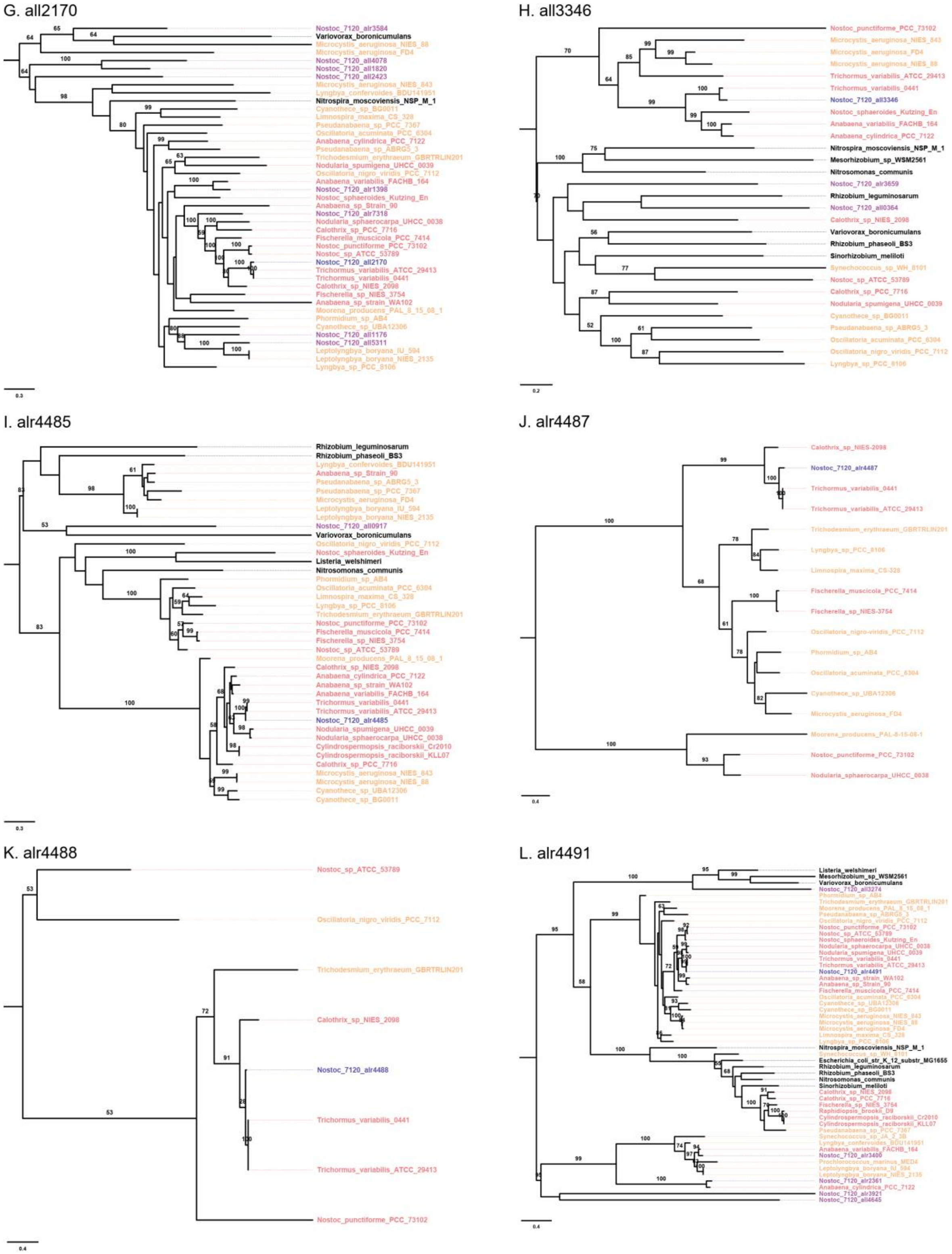
G. all2170
H. all3346
I. alr4485
J. alr4487
K. alr4488
L. alr4491

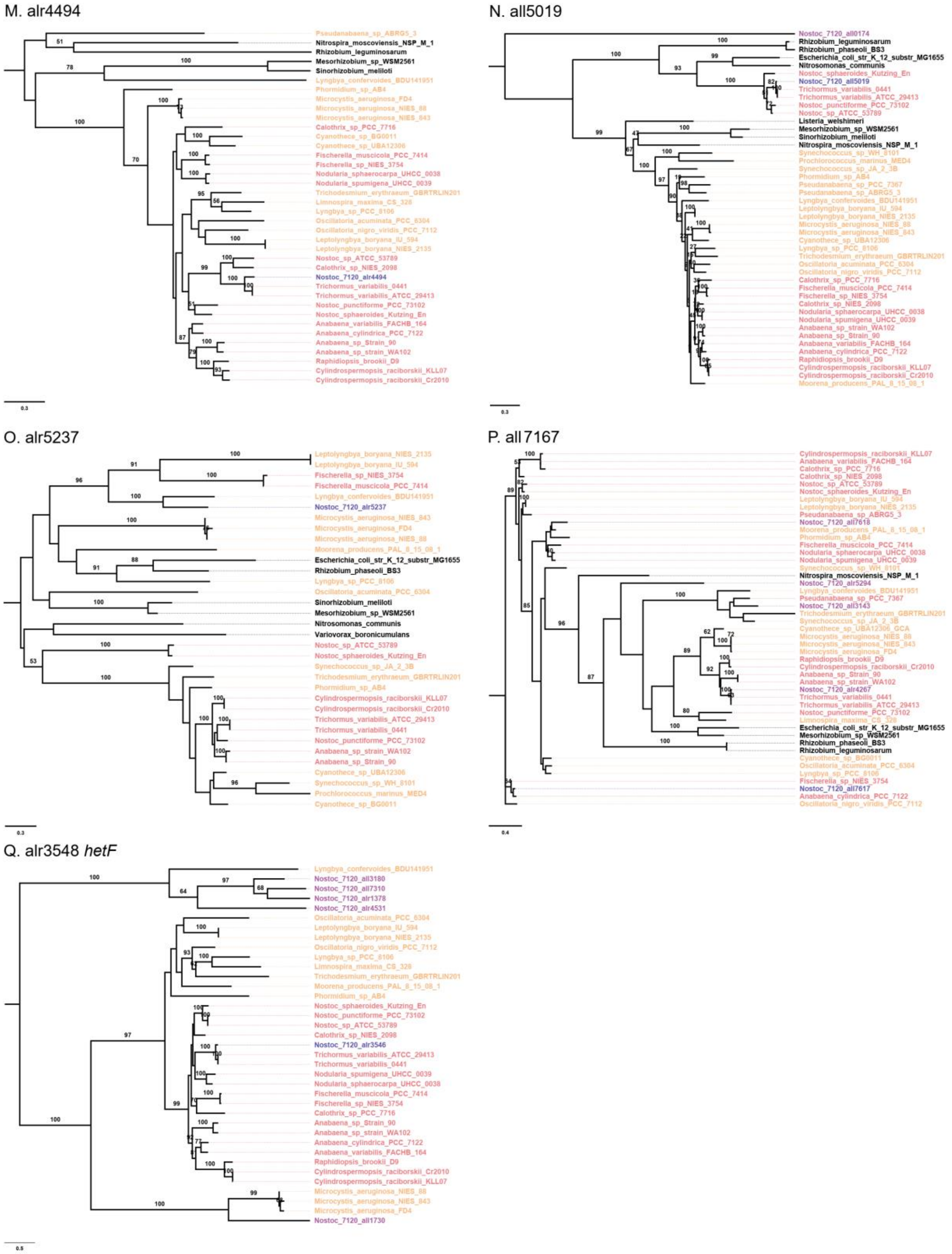


**Figure S4: Phylogeny of mutant genes.** Maximum likelihood trees of genes involved in resistance to phages and/or in nitrogen fixation in *Nostoc* 7120. The locus tag of the analyzed gene is marked above each phylogenetic tree. Blue, analyzed gene; purple, paralogs in *Nostoc* 7120; pink, homologs in other *Nostocales*; orange, homologs in non-*Nostocales cyanobacteria*; black, homologs in bacteria from different phyla. The database used for this analysis include genomes of 48 bacteria (Table S8).

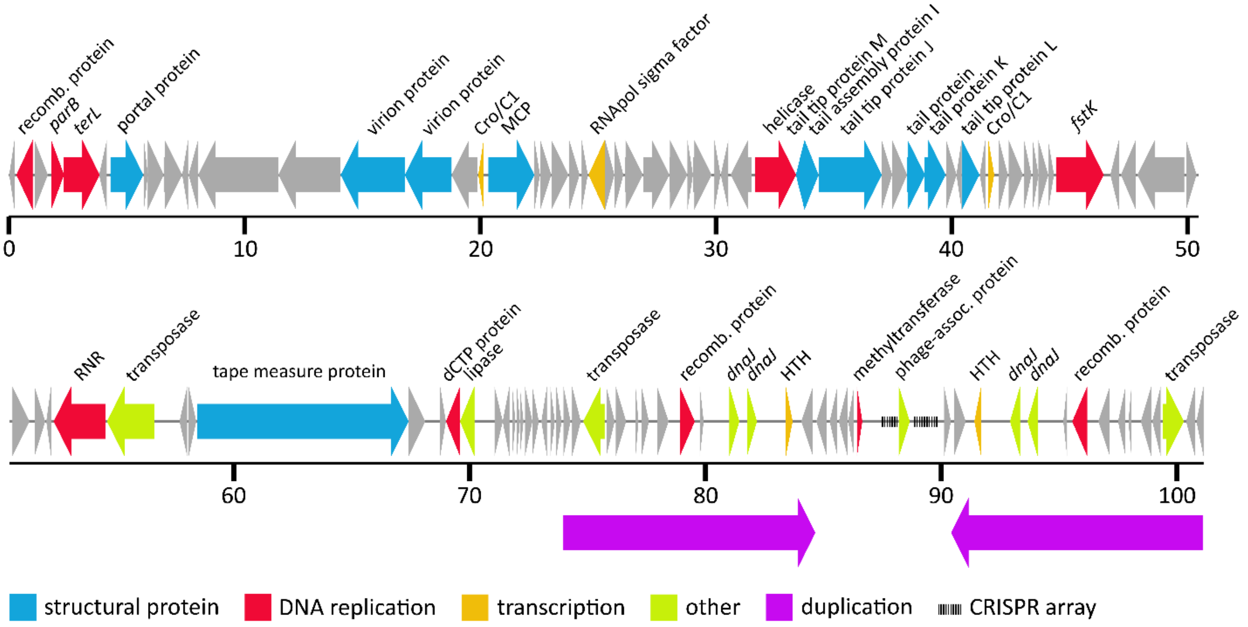


**Figure S5: Genome of Cylindrospermopsis phage Cr-LKS4.** Inverted duplication region is marked with purple arrows below the genome illustration. Recomb., recombination; MCP, major capsid protein; RNApol, RNA polymerase; RNR, ribonucleoside triphosphate reductase; dCTP, deoxycytidine triphosphate deaminase; HTH, helix-turn-helix domain-containing protein; assoc., associated.

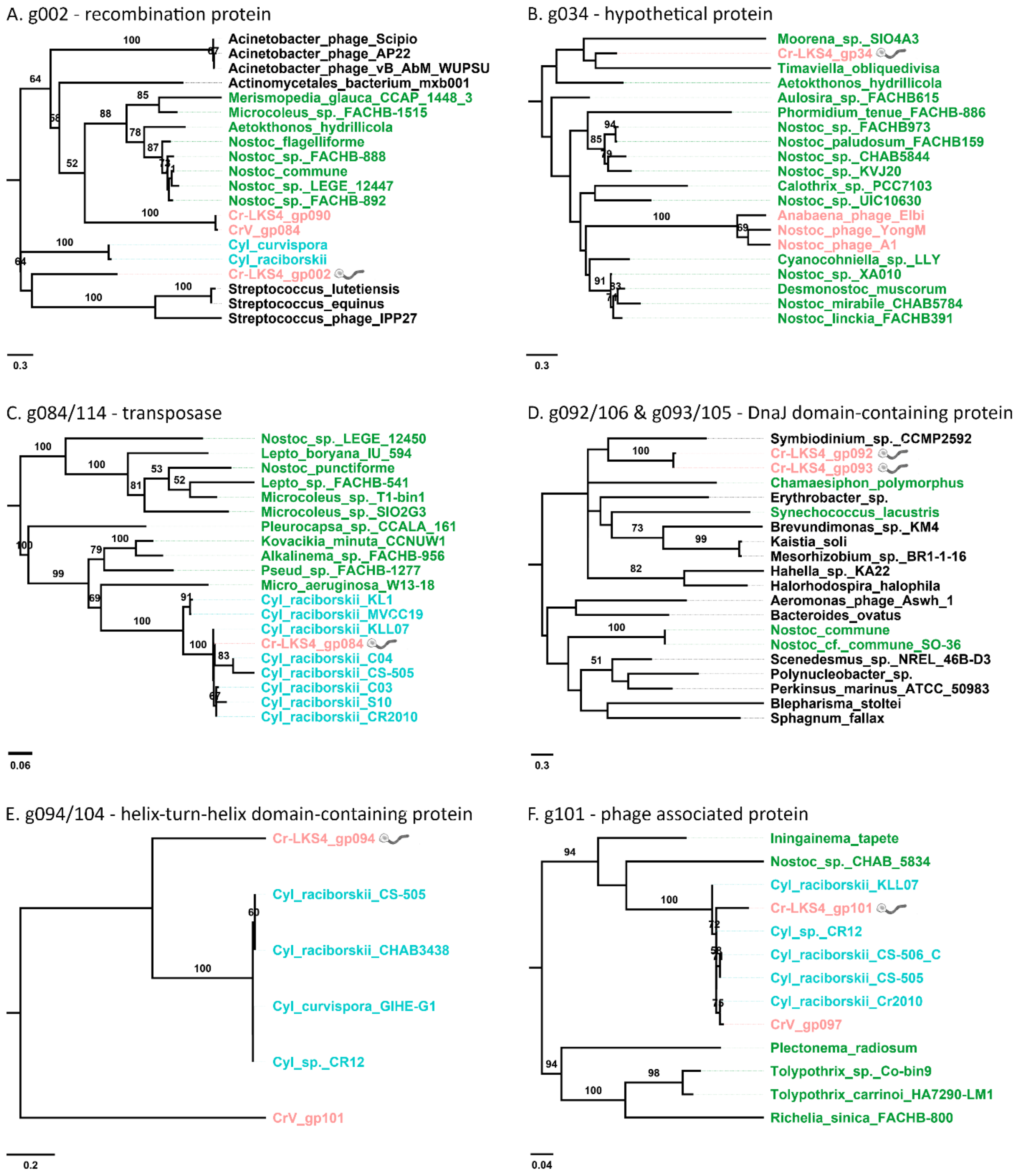


**Figure S6: Phylogeny of genes in Cylindrospermopsis phage Cr-LKS4.** Maximum likelihood trees of genes with a blastp best hit that was not in phage CrV. Light blue – *Cylindrospermopsis*, Green – other *cyanobacteria*, pink – cyanophage, black – other. The gene of interest in Cr-LKS4 is marked by a phage image. Bootstraps values greater than 50 (out of 100) are shown. Cyl, *Cylindrospermopsis*; Lepto, *Leptolyngbya*; Pseud, *Pseudomonas*; Micro, *Microcystis*.

#### References


1. Porankiewicz J, Wang J, Clarke AK. New insights into the ATP-dependent Clp protease: *Escherichia coli* and beyond. *Mol Microbiol* 1999; **32**: 449–458.

2. Nicolaisen K, Hahn A, Schleiff E. The cell wall in heterocyst formation by *Anabaena* sp. PCC 7120. *J Basic Microbiol* 2009; **49**: 5–24.

3. Xu X, Khudyakov I, Wolk CP. Lipopolysaccharide dependence of cyanophage sensitivity and aerobic nitrogen fixation in *Anabaena* sp. strain PCC 7120. *J Bacteriol* 1997; **179**: 2884–2891.

4. Kawano Y, Sekine M, Ihara M. Identification and characterization of UDP-glucose pyrophosphorylase in cyanobacteria *Anabaena* sp. PCC 7120. *J Biosci Bioeng* 2014; **117**: 531–538.

5. Katoh H, Asthana RK, Ohmori M. Gene Expression in the Cyanobacterium *Anabaena* sp. PCC7120 under Desiccation. *Microb Ecol* 2004; **47**: 164–174.

6. Curatti L, Giarrocco LE, Cumino AC, Salerno GL. Sucrose synthase is involved in the conversion of sucrose to polysaccharides in filamentous nitrogen-fixing cyanobacteria. *Planta* 2008; **228**: 617–625.

7. Wong FCY, Meeks JC. The *hetF* gene product is essential to heterocyst differentiation and affects *hetR* function in the cyanobacterium *Nostoc punctiforme*. *J Bacteriol* 2001; **183**: 2654–2661.

8. Wolk CP, Fan Q, Zhou R, Huang G, Lechno-Yossef S, Kuritz T, et al. Paired cloning vectors for complementation of mutations in the cyanobacterium *Anabaena* sp. strain PCC 7120. *Arch Microbiol* 2007; **188**: 551–563.

9. Risser DD, Callahan SM. HetF and PatA control levels of HetR in *Anabaena* sp. strain PCC 7120. *J Bacteriol* 2008; **190**: 7645–7654.

10. Xing W-Y, Liu J, Wang Z-Q, Zhang J-Y, Zeng X, Yang Y, et al. HetF protein is a new divisome component in a filamentous and developmental cyanobacterium. *mBio* 2021; **12**: e01382-21.

11. Aravind L, Koonin E V. Classification of the caspase-hemoglobinase fold: Detection of new families and implications for the origin of the eukaryotic separins. *Proteins Struct Funct Genet* 2002; **46**: 355–367.

12. Knowles EJ, Castenholz RW. Effect of exogenous extracellular polysaccharides on the desiccation and freezing tolerance of rock-inhabiting phototrophic microorganisms. *FEMS Microbiol Ecol* 2008; **66**: 261–270.

13. Qian N, Stanley GA, Hahn-Hägerdal B, Rådström P. Purification and characterization of two phosphoglucomutases from *Lactococcus lactis* subsp. *lactis* and their regulation in maltose- and glucose-utilizing cells. *J Bacteriol* 1994; **176**: 5304–5311.

14. Qian N, Stanley GA, Bunte A, Rdstrm P. Product formation and phosphoglucomutase activities in *Lactococcus lactis*: cloning and characterization of a novel phosphoglucomutase gene. *Microbiology* 1997; **143**: 855–865.

15. Levander F, Andersson U, Rådström P. Physiological Role of β-Phosphoglucomutase in *Lactococcus lactis*. *Appl Environ Microbiol* 2001; **67**: 4546–4553.

16. Lahiri SD, Zhang G, Dunaway-Mariano D, Allen KN. Caught in the act: The structure of phosphorylated β-phosphoglucomutase from *Lactococcus lactis*. *Biochemistry* 2002; **41**: 8351–8359.

17. Zeng X, Zhang CC. The making of a heterocyst in cyanobacteria. *Annu Rev Microbiol* 2022; **76**: 597–618.

18. Flores E, Arévalo S, Burnat M. Cyanophycin and arginine metabolism in cyanobacteria. *Algal Res* 2019; **42**: 101577.

19. Nikaido H, Hall JA. Overview of bacterial ABC transporters. *Methods Enzymol* 1998; **292**: 3–20.

20. Cuthbertson L, Kimber MS, Whitfield C, Raetz CRH. Substrate binding by a bacterial ABC transporter involved in polysaccharide export. *Proc Natl Acad Sci U S A* 2007.

21. Khudyakov I, Wolk CP. *hetC* , a gene coding for a protein similar to bacterial ABC protein exporters , is involved in early regulation of heterocyst differentiation in *Anabaena*

sp . strain PCC 7120. *J Bacteriol* 1997; **179**: 6971–6978.

22. Yen MR, Peabody CR, Partovi SM, Zhai Y, Tseng YH, Saier MH. Protein-translocating outer membrane porins of Gram-negative bacteria. *Biochim Biophys Acta - Biomembr* 2002; **1562**: 6–31.

23. Ngo G, Girbas M, Schätzle H, Hammer A, Safarian S, Hübinger M, et al. The two TpsB-Like proteins in *Anabaena* sp. strain PCC 7120 are involved in secretion of selected substrates. *J Bacteriol* 2021; **203**.

24. Forlani G, Bertazzini M, Barillaro D, Rippka R. Divergent properties and phylogeny of cyanobacterial 5-enol-pyruvyl-shikimate-3-phosphate synthases: Evidence for horizontal gene transfer in the Nostocales. *New Phytol* 2015; **205**: 160–171.

25. Flaherty BL, Van Nieuwerburgh F, Head SR, Golden JW. Directional RNA deep sequencing sheds new light on the transcriptional response of *Anabaena* sp. strain PCC 7120 to combined-nitrogen deprivation. *BMC Genomics* 2011; **12**: 332.

26. Cortazzo P, Cerveñansky C, Marín M, Reiss C, Ehrlich R, Deana A. Silent mutations affect in vivo protein folding in *Escherichia coli*. *Biochem Biophys Res Commun* 2002; **293**: 537–541.

27. Larkin RM, Alonso JM, Ecker JR, Chory J. GUN4, a regulator of chlorophyll synthesis and intracellular signaling. *Science* 2003; **299**: 902–906.

28. Davison PA, Schubert HL, Reid JD, Iorg CD, Heroux A, Hill CP, et al. Structural and biochemical characterization of gun4 suggests a mechanism for its role in chlorophyll biosynthesis. *Biochemistry* 2005; **44**: 7603–7612.

29. Formighieri C, Ceol M, Bonente G, Rochaix JD, Bassi R. Retrograde signaling and photoprotection in a gun4 mutant of *Chlamydomonas reinhardtii*. *Mol Plant* 2012; **5**: 1242–1262.

30. Bonente G, Formighieri C, Mantelli M, Catalanotti C, Giuliano G, Morosinotto T, et al. Mutagenesis and phenotypic selection as a strategy toward domestication of *Chlamydomonas reinhardtii* strains for improved performance in photobioreactors. *Photosynth Res* 2011; **108**: 107–120.

31. Pickard DJJ. Preparation of bacteriophage lysates and pure DNA. *Methods Mol Biol* 2009; **502**: 3–9.

32. Andrews S. FastQC: a quality control tool for high throughput sequence. http://www.bioinformatics.babraham.ac.uk/projects/fastqc/. 2010.

33. Nurk S, Bankevich A, Antipov D, Gurevich AA, Korobeynikov A, Lapidus A, et al. Assembling single-cell genomes and mini-metagenomes from chimeric MDA products. *J Comput Biol* 2013; **20**: 714–737.

34. Martin RM, Moniruzzaman M, Mucci NC, Willis A, Woodhouse JN, Xian Y, et al. *Cylindrospermopsis raciborskii* Virus and host: genomic characterization and ecological relevance. *Environ Microbiol* 2019; **21**: 1942–1956.

35. Besemer J. GeneMarkS: a self-training method for prediction of gene starts in microbial genomes. Implications for finding sequence motifs in regulatory regions. *Nucleic Acids Res* 2001; **29**: 2607–2618.

36. Wheeler DL, Church DM, Lash AE, Leipe DD, Madden TL, Pontius JU, et al. Database resources of the National Center for Biotechnology Information: 2002 update. *Nucleic Acids Res* 2002; **30**: 13–16.

37. Zimmermann L, Stephens A, Nam S-Z, Rau D, Kübler J, Lozajic M, et al. A completely reimplemented MPI Bioinformatics Toolkit with a new HHpred server at its core. *J Mol Biol* 2018; **430**: 2237–2243.

38. Couvin D, Bernheim A, Toffano-Nioche C, Touchon M, Michalik J, Néron B, et al. CRISPRCasFinder, an update of CRISRFinder, includes a portable version, enhanced performance and integrates search for Cas proteins. *Nucleic Acids Res* 2018; **46**: W246–W251.

39. Wheeler DL, Church DM, Lash AE, Leipe DD, Madden TL, Pontius JU, et al. Database resources of the National Center for Biotechnology Information: 2002 update. *Nucleic Acids Res* 2002; **30**: 13–16.

40. Katoh K, Rozewicki J, Yamada KD. MAFFT online service: Multiple sequence alignment, interactive sequence choice and visualization. *Brief Bioinform* 2018; **20**: 1160–1166.

41. Miller MA, Pfeiffer W, Schwartz T. Creating the CIPRES Science Gateway for inference of large phylogenetic trees. *2010 Gatew Comput Environ Work* 2010. IEEE, pp 1–8.
